# Global identification of *Chromobacterium violaceum* T6SS effectors reveals an Rhs antibacterial toxin featuring FIX and ADP-ribosyltransferase domains

**DOI:** 10.64898/2025.12.06.692724

**Authors:** Júlia A Alves, Genady Pankov, Andrew M Frey, Matthias Trost, Germán G Sgro, Sarah J Coulthurst, José F da Silva Neto

## Abstract

Bacteria coexist in polymicrobial communities where they engage in complex interactions, including interbacterial antagonism. We recently reported that the environmental bacterial pathogen *Chromobacterium violaceum* has an active type VI secretion system (T6SS), which plays a crucial role in interbacterial competition. However, the arsenal of toxic effectors delivered by this T6SS remains unknown. Here, we identify the repertoire of *C. violaceum* T6SS effectors and characterize a novel antibacterial Rhs-family effector, RhsF (Rhs with a FIX domain), and its cognate immunity protein, RhsFi. Using mass spectrometry analyses of secreted proteins and proteins co-immunoprecipitated with VgrG3, we identified six novel effector candidates, namely four phospholipases, a protein of unknown function, and the previously-uncharacterized Rhs protein, RhsF (CV_1431). RhsF contains an N-terminal FIX domain and was shown to intoxicate susceptible bacteria in a T6SS-dependent manner. The action of the C-terminal toxin domain of RhsF (RhsF-CT) is prevented in the presence of RhsFi (CV_1430), confirming that RhsF-RhsFi comprises an effector-immunity pair. The structure of the RhsF-CT/RhsFi complex determined by X-ray crystallography (1.85 Å resolution) revealed that RhsF-CT shares structural similarity with ADP-ribosyltransferase toxins and that RhsFi inhibits toxicity via direct occlusion of the RhsF-CT catalytic site. Functional assays confirmed that RhsF toxicity requires a catalytic triad composed of R1403, Y1456, and E1497 residues. Overall, our findings reveal the effectors secreted by the T6SS of *C. violaceum*, establish RhsF as a potent antibacterial toxin, and confirm T6SS-dependent delivery of a FIX-containing Rhs protein, expanding the known repertoire of bacterial arms involved in microbial competition.

## Introduction

Bacteria live in polymicrobial communities where a variety of complex ecological interactions takes place, including interbacterial competition. In this context, bacteria compete for nutrients and engage in interference competition, using diffusible molecules and toxins delivered by specialized secretion systems in order to inhibit or kill competitors (Peterson et al., 2020; Booth et al., 2023). The Type VI Secretion System (T6SS) is a contractile nanomachine employed by many Gram-negative bacteria for interbacterial competition, by delivering toxic effector proteins directly into neighbouring cells (Leiman et al., 2009; Zoued et al., 2014; Coulthurst, 2019). Among the four T6SS subtypes (T6SS^i^ to T6SS^iv^), the canonical T6SS^i^, predominantly found in Proteobacteria, is composed of three subcomplexes: a membrane complex (TssJLM), a baseplate (TssEFGK), and a contractile tail (TssBC) around an inner tube (Hcp) tipped by a cell-puncturing spike (VgrG-PAAR). Firing event involves contraction of the TssBC sheath, which propels the Hcp-VgrG-PAAR puncturing structure out of the secreting cell into the recipient cell. The Hcp-VgrG-PAAR puncturing structure is decorated with effector proteins, thereby translocating them into recipient/target cells (Basler et al., 2012; Böck et al., 2017; Unni et al., 2022). Anti-bacterial effectors will cause death or inhibition of growth of target bacterial cells unless they possess a cognate immunity protein. Effectors may be toxic C-terminal domains fused to the structural components VgrG, PAAR or Hcp (specialized effectors), or separate proteins that associate with VgrG, PAAR or Hcp non-covalently, sometimes via adaptor proteins (cargo effectors). Numerous T6SS effectors from distinct bacteria have been identified that act against competitor bacteria, by degrading the cell wall (amidases and glycosidases), the inner membrane (phospholipases), the nucleic acids (nucleases), or acting against other cellular targets (Coulthurst, 2019; Hernandez et al., 2020; Jurėnas and Journet, 2021). However, the repertoire of T6SS effectors remains to be experimentally determined for most bacterial species.

Rearrangement hot spot (Rhs) proteins are polymorphic toxins with a modular tripartite architecture (Zhang et al., 2012; Jamet and Nassif, 2015) that belong to the family of tyrosine/aspartate repeat-rich (YD-repeats) proteins found in bacteria, archaea, and eukaryotes (Tucker et al., 2012; Koskiniemi et al., 2013; Makarova et al., 2019). The N-terminal domain of Rhs proteins dictates their secretion pathway. In the case of T6SS-dependent secretion, many Rhs are specialised effectors containing an N-terminal PAAR domain (Cianfanelli et al., 2016; Jurėnas et al., 2021; Günther et al., 2022) or less commonly are fused to VgrG (Kielkopf et al., 2024). One exception is TseI from *Aeromonas dhakensis*, an Rhs with an N-terminal unknown function domain (Pei et al., 2020). The central core region of Rhs proteins is rich in tyrosine/aspartate (YD) repeats, which form a cocoon-like structure that protects the highly variable C-terminal toxic domain, delimited by the conserved PxxxxDPxGL motif (Pei et al., 2020; Jurėnas et al., 2021; Günther et al., 2022). Biochemical and structural studies indicate that the toxic activity of the C-terminal domain depends on its release from within the Rhs shell, where it is otherwise protected (Pei et al., 2020; Jurėnas et al., 2021; Günther et al., 2022; Tang et al., 2022; González-Magaña et al., 2023; Hayes et al., 2024). For instance, in *A. dhakensis*, the TseI effector undergoes autocleavage at both N- and C-terminal regions prior its interaction with VgrG for secretion (Pei et al., 2020). In *Photorhabdus laumondii*, Rhs1 contains N- and C-terminal plugs that seal the toxic Tre23 domain within the hollow barrel of the core shell (Jurėnas et al., 2021). In *Vibrio parahaemolyticus* RhsP, autoproteolysis induces significant structural rearrangements that facilitate β-barrel opening and RhsP dimerization (Tang et al., 2022). Like other T6SS effectors, the toxicity of Rhs proteins is neutralized by their cognate immunity proteins (Jamet and Nassif, 2015; Jurėnas and Journet, 2020; Pei et al., 2020; Bullen et al., 2022; Tang et al., 2022), yet structural information on Rhs effector-immunity complexes is still scarce.

*Chromobacterium violaceum* is a Gram-negative β-proteobacterium that inhabits soil and water in tropical and subtropical regions and causes rare but deadly infections in humans and other animals (de Vasconcelos et al., 2003; Miki et al., 2010; Batista and da Silva Neto, 2017). *C. violaceum* kills Gram-positive bacteria by delivering the purple antibiotic violacein into outer membrane vesicles (Durán et al., 2007; Batista et al., 2020). Recently, we demonstrated that *C. violaceum* strain ATCC 12472 outcompetes distinct Gram-negative bacteria using a single quorum sensing-regulated T6SS (Alves et al., 2022). We predicted potential T6SS effectors encoded by genes located in a main cluster and in four small orphan VgrG clusters in its genome (Alves et al., 2022). Here, we aimed to identify the antibacterial effectors secreted by the T6SS of *C. violaceum* through mass spectrometry analyses of secreted proteins in culture supernatants and proteins co-immunoprecipitated from cellular fractions with VgrG3, the most important among the six *C. violaceum* VgrG proteins. Combining these two approaches, we identified several novel potential effectors, adaptor proteins, all six VgrGs, and two PAAR proteins encoded outside of predicted T6SS clusters. Among the candidate T6SS-delivered effectors, we performed genetic, biochemical, and structural characterization of a new Rhs toxin-immunity pair, RhsF-RhsFi. Our data reveal that RhsF acts as an antibacterial toxin whose C-terminal toxic domain is inhibited by its cognate antitoxin, RhsFi, and whose toxic activity depends on a catalytic site resembling those of ADP-ribosyltransferase toxins. RhsF also represents the first experimentally-confirmed example of a T6SS-dependent Rhs effector with a FIX-containing N-terminal delivery domain.

## Results

### Global identification of effectors secreted by the *C. violaceum* T6SS

We previously demonstrated that the *C. violaceum* T6SS plays an important role in interbacterial competition and that VgrG3 is the most important among the six *C. violaceum*VgrG proteins for this T6SS-mediated killing (Alves et al., 2022). However, the T6SS-delivered antibacterial effectors remained to be identified. To identify effectors secreted by the *C. violaceum* T6SS, we employed two complementary strategies. The first involved using quantitative label-free mass spectrometry to compare the proteins in the secreted fraction (culture supernatant) of the wild type and the Δ*tssB* mutant (in which T6SS secretion is inactivated). This resulted in the identification of 10 non-redundant proteins significantly enriched, or exclusively present, in supernatant of the wild type strain, indicating that they are secreted in a T6SS-dependent manner (Figs. 1A, C, S1A, Table 1). Of these, five proteins are secreted T6SS core components (Hcp, VgrG2, VgrG3, VgrG4, and VgrG6), two are phospholipases of the Tle1 family (CV_3990, associated with VgrG1, and CV_3971, associated with VgrG2), and one is a hypothetical protein (CV_2125), whose gene is organized in operon with a gene encoding a PAAR protein (Fig. 1D). The high similarity among the six VgrGs encoded by this bacterium, which share 71 to 93% amino acid identity, makes it challenging to identify specific VgrGs by mass spectrometry analysis. For that reason, some hits do not correspond to a specific VgrG (Fig. 1A, Table 1).

**Figure 1.**
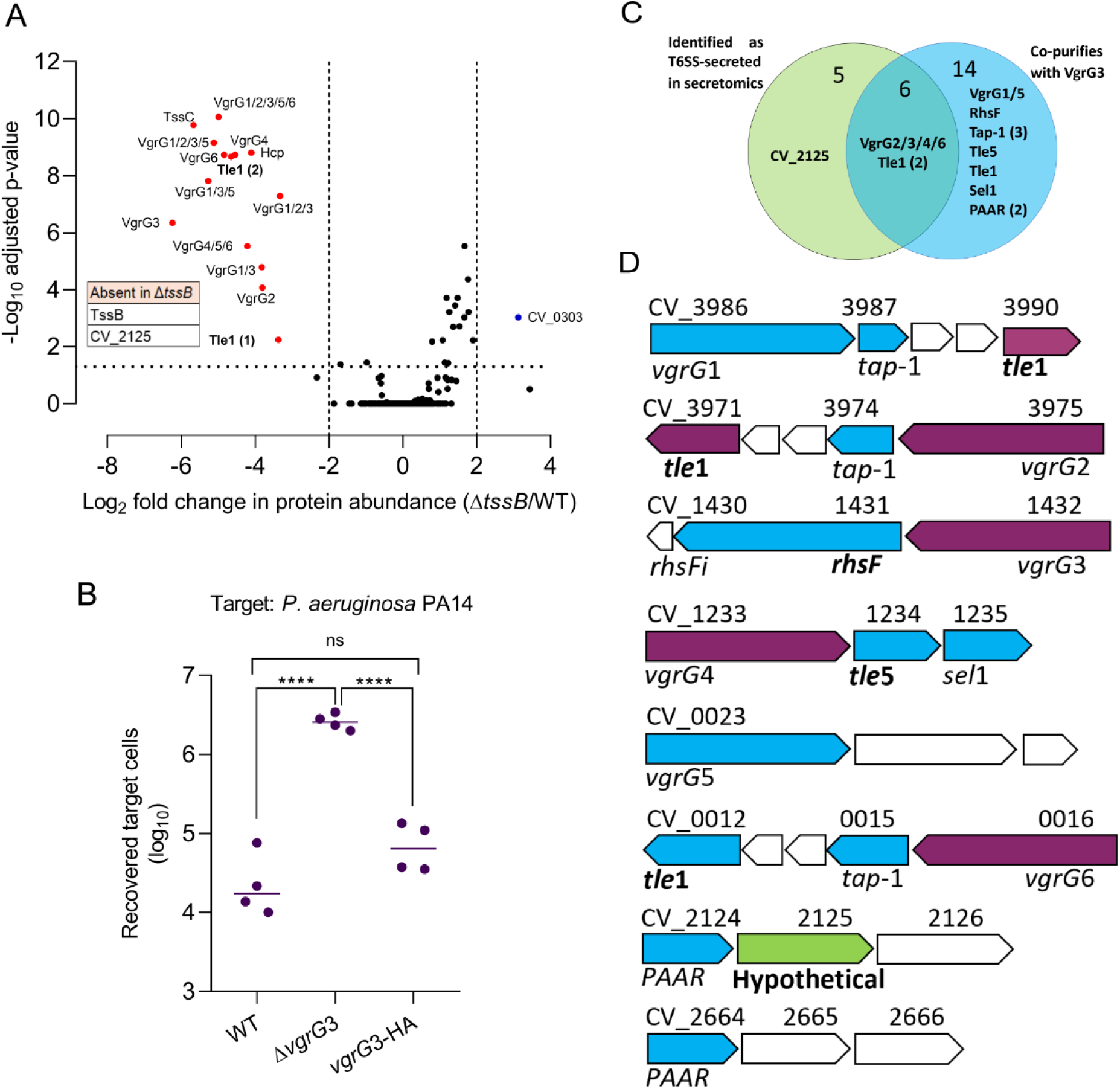
Global identification of proteins secreted by the T6SS of *C. violaceum*. *A,* label-free quantitative mass spectrometry analysis of the secreted protein fraction obtained from *C. violaceum* grown in minimal medium. The volcano plot summarizes the comparison of proteins identified in the culture supernatant of wild-type *C. violaceum* ATCC 12472 (WT) and the Δ*tssB* mutant. Colored dots indicate proteins with significantly altered abundance in the Δ*tssB* mutant compared with the wild type (log2 fold change ≤ -2 or ≥ 2, *p*-value < 0.05, n=7 independent biological replicates). Proteins CV_2125 and TssB were identified in the wild type supernatant samples but were not detected in Δ*tssB* and therefore are not plotted. Further details for individual proteins are given in Table 2. *B,* inter-species T6SS-dependent antibacterial activity. Recovery of *P. aeruginosa* PA14 target cells following co-culture with wild type *C. violaceum* or mutants carrying an in-frame deletion of *vgrG3* (Δ*vgrG3*) or encoding VgrG with a C-terminal HA tag at the normal chromosomal location (*vgrG*3-HA). Data are presented with a line showing the mean and individual data points overlaid (n=4 biological replicates; **** P<0.0001, ns not significant; one-way ANOVA with Tukey’s test). *C,* Venn diagram showing the number of proteins identified by each strategy, with proteins of interest highlighted. *D*, genomic organization of genes encoding the identified T6SS-associated proteins. Blue arrows: proteins identified via VgrG3 co-immunoprecipitation (Co-IP); Green arrows: proteins identified via secretomics analysis; Purple arrows: proteins identified by both approaches. Genes encoding potential effectors are indicated in bold.

**Table 1.** T6SS-dependent proteins of *C. violaceum* identified by label-free quantitative mass spectrometry analysis of the secretome of the wild type and Δ*tssB* mutant.

| Accession | Identifier | Description | Log10 adjusted *P*-value | Log2 fold change $\Delta tssB$/WT | Number of peptides |
| --- | --- | --- | --- | --- | --- |
| Q7NR06 | CV_3979 | TssB (T6SS sheath protein) | ND | ND | 4 |
| Q7NW65 | CV_2125 | Uncharacterized protein | ND | ND | 5 |
| Q7P238;<br>Q7P245 | CV_0016;<br>CV_0023 | VgrG5/6 | ND | ND | 3 |
| Q7NY43 | CV_1432 | VgrG3 | 6.34 | -6.24 | 6 |
| Q7NR07 | CV_3978 | TssC (T6SS sheath protein) | 9.775 | -5.67 | 19 |
| Q7NQZ9;<br>Q7NY43;<br>Q7P238 | CV_0023;<br>CV_1432;<br>CV_3986 | VgrG1/3/5 | 7.815 | -5.26 | 3 |
| Q7NQZ9;<br>Q7NR10;<br>Q7NY43;<br>Q7P238 | CV_0023;<br>CV_1432;<br>CV_3975;<br>CV_3986 | VgrG1/2/3/5 | 9.153 | -5.12 | 4 |
| Q7NQZ9;<br>Q7NR10;<br>Q7NY43;<br>Q7P238;<br>Q7P245 | CV_0016;<br>CV_0023;<br>CV_1432;<br>CV_3975;<br>CV_3986 | VgrG1/2/3/5/6 | 10.057 | -4.99 | 6 |
| Q7P245 | CV_0016 | VgrG6 | 8.727 | -4.84 | 4 |
| Q7NR10 | CV_3975 | VgrG2 | 4.078 | -3.81 | 12 |
| Q7NR14 | CV_3971 | Tle1 (phospholipase effector) | 8.665 | -4.65 | 33 |
| Q7NYP0 | CV_1233 | VgrG4 | 8.727 | -4.54 | 21 |
| Q7NYP0;<br>Q7P238;<br>Q7P245 | CV_0016;<br>CV_0023;<br>CV_1233 | VgrG4/5/6 | 5.525 | -4.21 | 5 |
| Q7NQZ9;<br>Q7NY43 | CV_1432;<br>CV_3986 | VgrG1/3 | 4.784 | -3.82 | 6 |
| Q7NR08 | CV_3977 | Hcp | 8.806 | -4.1 | 29 |
| Q7NQZ5 | CV_3990 | Tle1 (phospholipase effector) | 2.241 | -3.37 | 13 |
| Q7NQZ9;<br>Q7NR10;<br>Q7NY43 | CV_1432;<br>CV_3975;<br>CV_3986 | VgrG1/2/3/5 | 7.286 | -3.32 | 10 |
| Q7P1B0 | CV_0303 | Probable indolepyruvate ferredoxin oxidoreductase | 3.029 | 3.13 | 45 |

Proteins are included on the basis of the their abundance in the secreted fraction being > 4-fold altered in the $\Delta tssB$ mutant compared with the wild type (log2 fold change $\leq -2$ or $\geq 2$ ) with a significant t-test ( $p < 0.05$ ), or if they were not identified in the secreted fraction of $\Delta tssB$ but were detected in all replicates of the wild type. The table summarises the data from seven independent biological replicates of each strain. ND stands for not detected.

**Table 2.** VgrG3-associated proteins identified by co-immunoprecipitation.

| Accession | Gene ID | Description | Mean LFQ intensity VgrG3-HA | Mean LFQ intensity WT | Peptides | Unique peptides | Sequence coverage [%] | -Log10 p-val VgrG3-HA/WT | Log2 FC VgrG3-HA/WT |
| --- | --- | --- | --- | --- | --- | --- | --- | --- | --- |
| Q7NQZ9 | CV_3986 | VgrG1 | 31.23 | ND | 102 | 11 | 89.7 | ND | ND |
| Q7NR11 | CV_3974 | DUF4123 domain-containing protein Tap-1 | 31.24 | ND | 25 | 25 | 79 | ND | ND |
| Q7NTA8 | CV_3151 | Uncharacterized protein | 27.47 | ND | 17 | 17 | 45.3 | ND | ND |
| Q7NUN3 | CV_2664 | PAAR domain-containing protein | 26.95 | ND | 9 | 9 | 62.6 | ND | ND |
| Q7NW66 | CV_2124 | PAAR domain-containing protein | 29.04 | ND | 14 | 14 | 93.3 | ND | ND |
| Q7NYN8 | CV_1235 | Lipoprotein Sel-1 | 31.34 | ND | 33 | 33 | 72.1 | ND | ND |
| Q7NZ98 | CV_1024 | Flagellar biosynthesis protein FlhF | 27.27 | ND | 16 | 16 | 57.2 | ND | ND |
| Q7P246 | CV_0015 | DUF4123 domain-containing protein Tap-1 | 29.84 | ND | 21 | 21 | 64.5 | ND | ND |
| Q7NQZ8 | CV_3987 | DUF4123 domain-containing protein Tap-1 | 31.66 | 24.90 | 25 | 25 | 85.9 | ND | 6.76 |
| Q7P245 | CV_0016 | VgrG6 | 34.07 | 26.67 | 95 | 41 | 90.8 | ND | 7.40 |
| Q7P238 | CV_0023 | VgrG5 | 33.20 | 22.74 | 101 | 17 | 88.2 | 4.93 | 10.46 |
| Q7NR14 | CV_3971 | Tle1 phospholipase | 32.71 | 23.14 | 67 | 67 | 76 | 4.35 | 9.57 |
| Q7NY44 | CV_1431 | RhsF | 36.69 | 27.54 | 241 | 241 | 95.8 | 9.75 | 9.15 |
| Q7NQZ5 | CV_3990 | Tle1 phospholipase | 32.38 | 24.10 | 44 | 44 | 60.8 | 3.63 | 8.28 |
| Q7NR10 | CV_3975 | VgrG2 | 32.87 | 25.53 | 97 | 38 | 89.6 | 2.40 | 7.34 |
| Q7P249 | CV_0012 | Tle1 phospholipase | 30.78 | 23.47 | 38 | 38 | 71.5 | 7.30 | 7.31 |
| Q7NYN9 | CV_1234 | Tle5 phospholipase | 31.62 | 24.33 | 54 | 54 | 86.3 | 5.33 | 7.29 |
| Q7NYP0 | CV_1233 | VgrG4 | 35.67 | 28.54 | 121 | 109 | 92.7 | 4.82 | 7.13 |
| Q7NY43 | CV_1432 | VgrG3 | 36.56 | 29.56 | 129 | 38 | 92.2 | 6.06 | 7.01 |
| Q7NYC6 | CV_1348 | Threonine--tRNA ligase. ThrS | 26.97 | 24.72 | 14 | 14 | 33.5 | 2.39 | 2.25 |

Only proteins identified in all three replicates of VgrG3-HA are included. Proteins identified in two or three replicates of the control (WT) were excluded if they had an LFQ intensity value < 25 or a log2 fold change (VgrG3-HA/WT) < 3. ND stands for not detected.

The second strategy used to identify new T6SS effectors was to identify proteins interacting with VgrG3 by co-immunoprecipitation from total cellular protein. For this purpose, a strain encoding a fusion of VgrG3 with a C-terminal haemagglutinin epitope tag (VgrG3-HA) at the normal chromosomal location was generated. The functionality of VgrG3-HA was confirmed by determining T6SS-dependent anti-bacterial activity against *Pseudomonas aeruginosa* in a co-culture assay, where the *vgrG3-HA* strain displayed similar activity to the wild type, in contrast with the Δ*vgrG3* mutant (Fig. 1B). Using quantitative mass spectrometry, 20 proteins were identified as being exclusively present, or significantly enriched, in the VgrG3-HA co-immunoprecipitation compared with the control (Table 2, Fig. S1B, C). Of these, six proteins were also identified as being present in the secretome in a T6SS-dependent manner: the Tle1 phospholipases CV_3990 and CV_3971 and the VgrG proteins VgrG2, VgrG3, VgrG4, and VgrG6 (Fig. 1C, D). An additional fourteen proteins were identified, including: VgrG1, VgrG5, another Tle1 family phospholipase (CV_0012), three Tap-1 adaptor proteins (CV_3987, CV_3974, and CV_0015), one Tle5 family phospholipase (CV_1234) and its putative immunity protein Sel-1 (CV_1235), one Rhs family protein (CV_1431), and two PAAR-containing proteins (CV_2124 and CV_2664) (Fig. 1C, D, Table 2). Taken together, the two strategies revealed many candidate *C. violaceum* T6SS effectors, including four phospholipases, a protein of unknown function (CV_2125), and an uncharacterized Rhs protein (CV_1431).

### RhsF is an antibacterial toxin delivered by the *C. violaceum* T6SS and neutralized by RhsFi

In the VgrG3 co-immunoprecipitation, we identified an Rhs family protein (CV_1431) that we named RhsF (for Rhs family protein containing a FIX domain). Analysis of this protein revealed a modular architecture typical of Rhs proteins (Pei et al., 2020; Jurėnas et al., 2021; Günther et al., 2022). RhsF contains a FIX domain, which is associated with a number of T6SS substrates (Jana et al., 2019), in its N-terminal portion, a central Rhs core repeat region, and a C-terminal domain delimited by the PxxxxDPxGL motif (Fig. 2A). The *rhsF* gene is organized in a putative operon with *vgrG*3 (CV_1432) and a gene encoding a small hypothetical protein (CV_1430) that we named RhsFi (cognate immunity protein of RhsF) (Fig. 1D). To assess whether the C-terminal domain of RhsF (RhsF-CT) displays antibacterial toxicity, a gene encoding RhsF-CT (from after the predicted cleavage site delineated by the PxxxxDPxGL motif to the end of the protein, amino acids 1393-1513) was cloned under the control of an L-arabinose-inducible promoter, with and without the gene encoding the predicted immunity protein, RhsFi. Using these constructs to assess the impact of RhsF-CT expression revealed that the RhsF-CT is toxic when expressed in *Escherichia coli* and that this toxicity is neutralized by co-expression with RhsFi (Figs. 2B, S2), indicating that RhsF-RhsFi comprise an effector-immunity pair.

**Figure 2.**
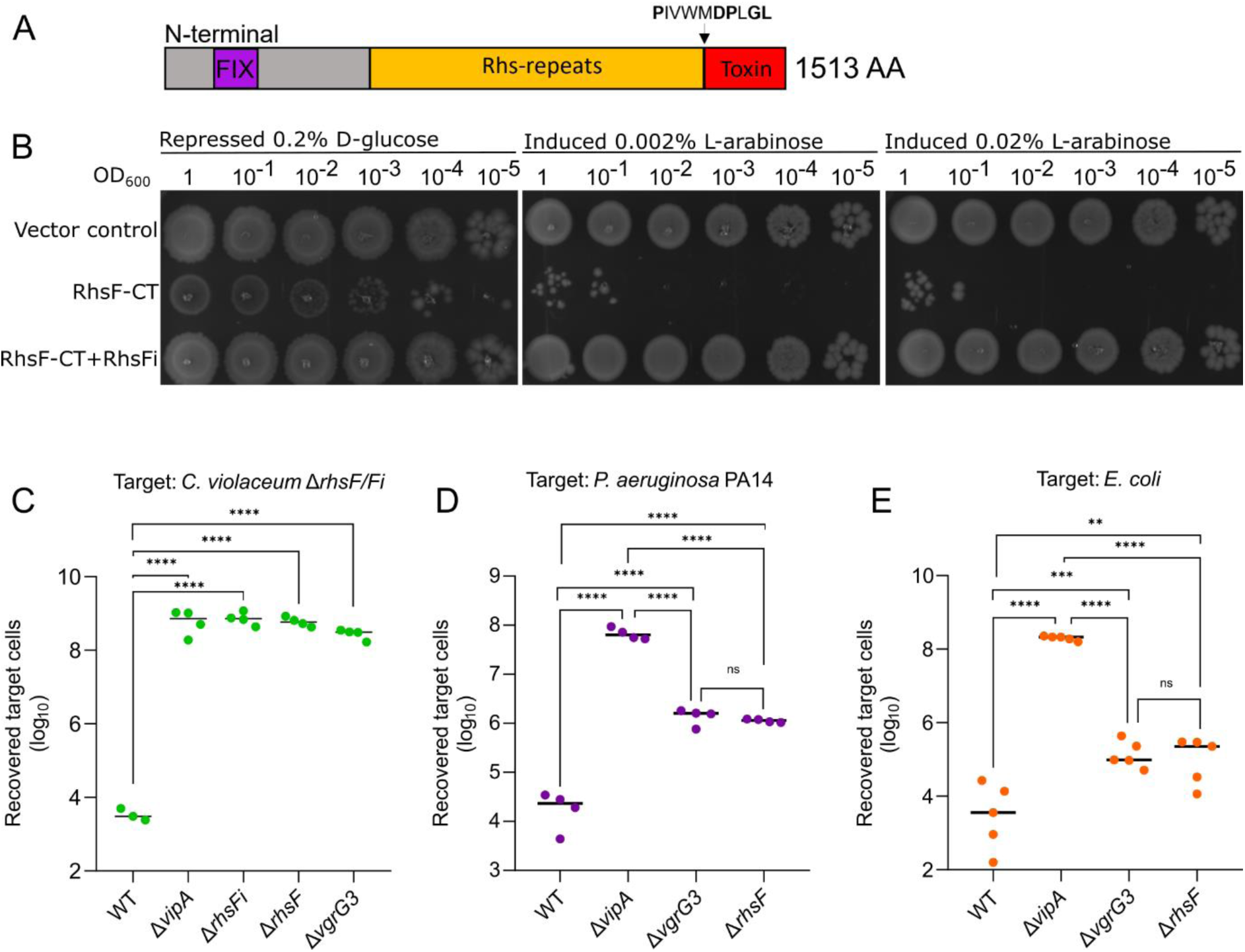
RhsF is an antibacterial effector delivered by the T6SS. *A,* modular organization of RhsF highlighting the N-terminal FIX domain, the central Rhs repeat core, and the C-terminal toxic domain delimited by the conserved PxxxxDPxGL motif. *B,* growth of *E. coli* MG1655 carrying pBAD18-Kan (vector control) or derivatives directing the expression of the C-terminal toxin domain of RhsF (RhsF-CT) alone or co-expressed with RhsFi on LB media containing either D-glucose (repression) or L-arabinose (induction) for regulation of gene expression. *C,* intraspecies T6SS-dependent antibacterial activity. Recovery of *C. violaceum* Δ*rhsF/Fi* (target) following co-culture with wild type *C. violaceum* or mutants carrying in-frame deletions. *D-E*, interspecies T6SS-dependent antibacterial activity. Recovery of *P. aeruginosa* PA14 and *E. coli* following co-culture with wild type *C. violaceum* or mutants carrying in-frame deletions. *C-E*, Data are presented with a line showing the mean and individual data points overlaid (n≥ 3 biological replicates; ****, P < 0.0001; ***, P<0.001; **, P < 0.01; ns not significant; one-way ANOVA with Tukey’s *post hoc* test).

To determine whether the secretion of RhsF is dependent on the T6SS, we performed a co-culture assay for T6SS-dependent antibacterial activity. In this assay, a mutant strain of *C. violaceum* lacking the immunity protein RhsFi (in-frame deletions of *rhsF* and *rhsFi,* Δ*rhsF/Fi*) and marked with nalidixic acid resistance was used as the ‘target’ strain. Recovery of this RhsF-susceptible target strain was determined following co-culture with strains of *C. violaceum* with or without the ability to deliver RhsF. The number of colony-forming units of Δ*rhsF/Fi* recovered following competition against the *C. violaceum* wild-type strain was approximately 4 logs lower than following competition with strains with an inactive T6SS (Δ*tssB*) or lacking *rhsF* or *vgrG*3 (Fig. 2C), indicating that the delivery of RhsF is dependent on both the T6SS and VgrG3. The T6SS is capable of secreting different effectors (toxins) in a single firing event, and this diversity ensures greater efficiency in killing competing bacteria (Smith et al., 2025). Given that we identified several proteins secreted by the *C. violaceum* T6SS that potentially act as antibacterial effectors (phospholipases from the Tle1 and Tle5 families, the hypothetical protein CV_2125, and RhsF), we evaluated the role of RhsF within this cocktail of secreted toxins. Interbacterial competition assays were conducted using *E. coli* and *P. aeruginosa* PA14 as targets. In both cases, deletion of *rhsF* significantly reduced the killing potential of *C. violaceum* compared to the wild-type strain (Fig. 2D, E), indicating that RhsF plays an important role in diversifying the toxins secreted by the T6SS, through the toxic action of its CT and/or supporting delivery of other effectors as part of the secreted T6SS machinery.

### Structure of the RhsF-CT/RhsFi complex provides insights into toxin inhibition

To investigate the possible mechanism of action of the RhsF toxin, we initially used the RhsF-CT amino acid sequence to search for related proteins using protein Basic Local Alignment Search Tool (BLASTp) and for conserved motifs using the Motif Finder tool. Since no significant matches were retrieved, we attempted to determine the structure of RhsF-CT. His6-tagged RhsF-CT (amino acids 1393 to 1513 of RhsF) was co-produced with its immunity protein, RhsFi, and a stable complex of the two proteins was isolated by nickel-affinity chromatography (Fig. S3A). This complex was then subjected to size exclusion chromatography, where it eluted as a single peak with an estimated molecular weight of ∼29 kDa (Fig. S3B). Since RhsF-CT has a molecular weight of 14 kDa and RhsFi of 12 kDa, these data suggest the formation of a stable, heterodimeric RhsF-CT/RhsFi complex with a 1:1 stoichiometry. The complex was subjected to crystallization trials and following optimization of crystallization conditions, diffracting crystals were obtained (Fig. S3C).

We determined a structure of RhsF-CT in complex with RhsFi to a resolution of 1.85 Å (Table 3; Fig. 3A, B). The complex adopts a globular conformation in which RhsF-CT and RhsFi interact in a 1:1 stoichiometric ratio (Fig. 3A). The RhsF-CT has a structural core composed of a β-sheet divided into two units of antiparallel strands (β1, β3, β5, and β2, β4) interspersed with α-helices (Fig. 3B), whereas the immunity protein RhsFi is composed of six α-helices interspersed with small loops (Fig. 3C). The protein:protein interface covers approximately 1007.7 Å^2^, representing 13.5% of the solvent-accessible area of RhsF-CT (total area 7584.1 Å^2^) and 15.8% of RhsFi (total area 6300.1 Å^2^). The interaction is mediated by 28 residues of RhsF (22.2% of total residues) and 26 residues of RhsFi (25% of total residues), that form 12 hydrogen bonds and 15 salt bridges (Figure S4).

**Figure 3.**
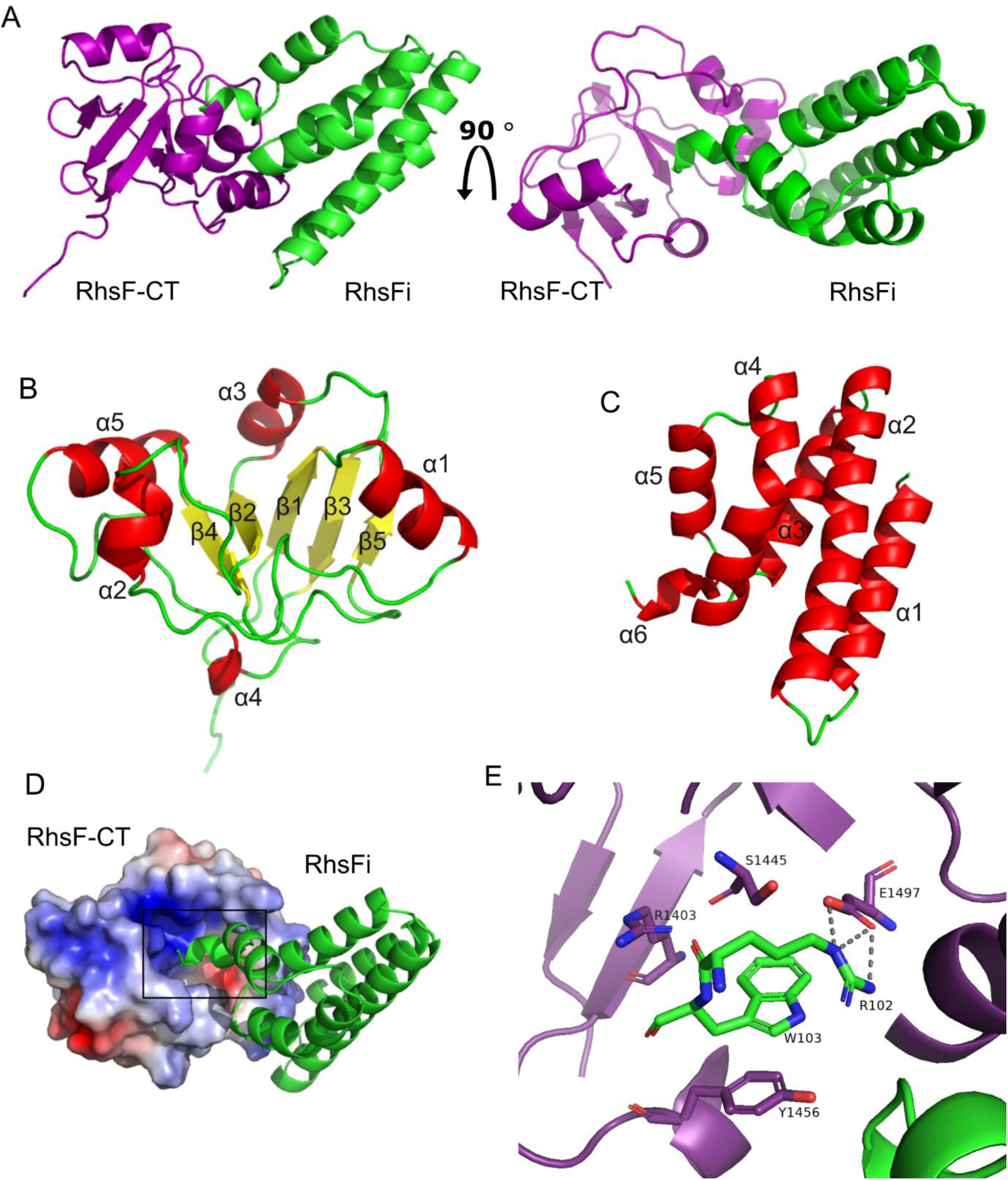
Crystal structure of the RhsF-CT/RhsFi complex. *A,* cartoon representation of the RhsF-CT/RhsFi complex. RhsF-CT is shown in purple and RhsFi in green. *B-C,* cartoon representations of RhsF-CT (*B*) and RhsFi (*C*), with α-helices in red, β-strands in yellow, and loops in green. *D,* representation of the RhsF-CT/RhsFi complex showing the surface electrostatic potential of RhsF-CT, with positively charged regions in blue and negatively charged regions in red. RhsFi is shown in green cartoon. Box highlight interaction of RhsFi with putative catalytic site. *E*, close-up view of part of the interaction interface between RhsF-CT (toxin) and RhsFi (immunity protein). Key residues at the interface are highlighted: E1497, R1403, S1445, and Y1456 in the toxin (RhsF-CT, purple), and R102 and W103 in the antitoxin (RhsFi, green).

**Table 3.** Data collection and structural refinement statistics.

| RhsF/RhsFi complex |  |
| --- | --- |
| **Data collection** |  |
| Space group | P1 |
| Cell Dimensions |  |
| a, b, c (Å) | 34.83, 52.92, 66.39 |
| α, β, γ (°) | 72.03, 86.91, 71.84 |
| Resolution range (Å) | 47.85 - 1.85 |
| *R*<sub>merge</sub> | 0.087 |
| *I*/*σI* | 4.34 (1.85) |
| Completeness (%) | 93.6 (47.85-1.85) |
| Redundancy | 3.4 |
| **Refinement** |  |
| Resolution (Å) | 1.85 |
| No. reflections | 34223 |
| *R*<sub>work</sub>/*R*<sub>free</sub> | 0.1899 / 0.2194 |
| No. atoms | 7752 |
| Protein | 7314 |
| Ligand/ion | 0 |
| Water | 438 |
| Average *B*-factors | 29.82 |
| Protein | 29.14 |
| Ligand/ion | 0 |
| Water | 35.48 |
| R.m.s. deviations |  |
| Bond lengths (Å) | 0.007 |
| Bond angles (°) | 0.85 |

Notably, the RhsF-CT features an electropositive pocket, its putative catalytic site, where four residues (R1403, S1445, Y1456, and E1497) interact closely with R102 and W103 in the α6 helix of RhsFi (Fig. 3D, E). In this interface, R102 of RhsFi engages E1497 of RhsF-CT through an ionic bridge, and W103 directly interacts with RhsF residues S1445, Y1456, and R1403 via π–π and cation–π interactions (Gallivan and Dougherty, 1999; Shao et al., 2022).

Immunity proteins associated with antibacterial effectors secreted by the T6SS typically function by occluding the catalytic site of the effector (Benz et al., 2012; Russell et al., 2014). Therefore, the residues of RhsF-CT that interact with RhsFi may be key to understanding the function of this toxin.

### RhsF-CT exhibits structural and catalytic features of ADP-ribosyltransferase toxins

A search for proteins with structural similarity to RhsF-CT was conducted using PDB e-Fold (Krissinel and Henrick, 2004). Three entries were retrieved using RMSD ≤ 2.5 and Z-score ≥ 5.0 as a threshold: the ScARP protein from *Streptomyces coelicolor*, bound and unbound with NADH (PDB 5zj5 and 5zj4, respectively), and the EcPltA protein from *E. coli* (PDB 4z9d) (Table S1). ScARP is an ADP-ribosyltransferase (ART) which ADP-ribosylates guanine nucleosides (Nakano et al., 2013; Yoshida and Tsuge, 2018). EcPltA is also an ADP-ribosyltransferase, encoded by certain pathogenic extraintestinal *E. coli* strains, which ADP-ribosylates mammalian inhibitory trimeric G-proteins, exhibiting cytotoxic activity (Littler et al., 2017). Bacterial ADP-ribosyltransferase toxins utilize NAD as a cofactor to catalyse the transfer of an ADP-ribose moiety onto an acceptor substrate, disrupting its function (Simon et al., 2014; Suskiewicz et al., 2023). There is significant sequence divergence among bacterial ARTs, which can hinder their characterization based solely on sequence analysis. However, ARTs share a conserved core structure composed of a β-sheet formed by three antiparallel strands interspersed with α-helices (Suskiewicz et al., 2023), a feature that is also present in RhsF-CT (Fig. 3B). Structural superimposition of RhsF-CT with ScARP and EcPltA confirmed a shared structural similarity among these proteins (Fig. 4A).

**Figure 4.**
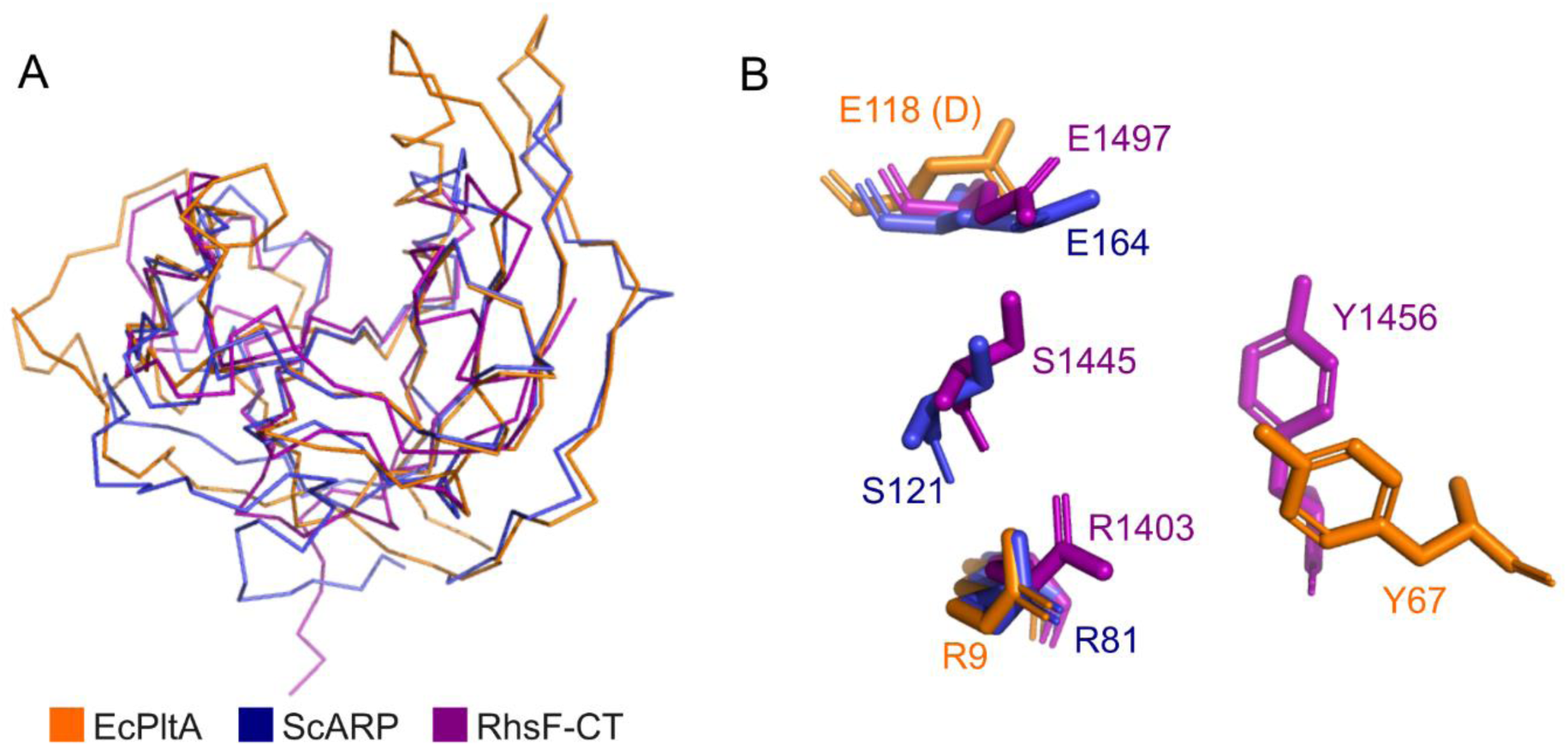
RhsF-CT shares structural similarity with ADP-ribosyltransferases. *A,* superimposition of RhsF-CT with ScARP (PDB 5zj4) and EcPltA (PDB 4z9d). Structures are depicted in ribbon representation, with EcPltA in orange, ScARP in blue and RhsF-CT in purple. *B,* close-up view of the relative positions of key residues from EcPltA and ScARP, along with their counterparts in RhsF-CT, based on the structural alignment in *A*. EcPltA residue E118 is shown as an aspartate (D) because the available structure contains this substitution.

The ART superfamily can be divided into two major groups depending on domain organization and conservation of active site motifs. Bacterial ARTs of class I, a group that includes diphtheria toxin, possess an active site composed of H-Y-E motif, in which the nucleophilic histidine is proposed to bind to nicotinamide group, orienting NAD molecule in the catalytic pocket. The invariable glutamic acid residue is the key catalytic residue responsible for coordination of NAD for hydrolysis prior to its binding to the substrate. ARTs of class II, on the other hand, present a triad composed of R-S-E motif, in which the nucleophilic arginine and serine residues positions and stabilizes the NAD molecule in the active site, and the highly conserved glutamic acid has a similar function as in class I ARTs (Domenighini et al., 1994, Simon et al., 2014; Suskiewicz et al., 2023). The catalytic residues in ScARP and EcPltA are R81-S121-E164 and R9-Y67-E118, respectively. We performed a structural superimposition to identify the equivalent residues in RhsF-CT. This indicated a high degree of spatial conservation with the arginine and glutamate residues from ScARP and EcPltA, which aligned with R1403 and E1497 in RhsF, respectively (Fig. 4B). The third component of the catalytic triad, however, could not be unambiguously identified by this method, as Y1456 in RhsF aligns with Y67 from EcPltA, while S1445 aligns with S121 from ScARP (Fig. 4B).

To test the importance of the predicted catalytic triad residues of RhsF, the toxicity of variants of RhsF-CT carrying substitutions of these residues was tested by heterologous expression in *E. coli*. The growth of cultures of *E. coli* MG1655 carrying constructs directing the expression of RhsF-CT variants was assessed in both solid and liquid LB medium (Fig. 5A, B). Individual substitution of residues E1497, R1403, and Y1456 with alanine resulted in loss of RhsF-CT toxicity in both conditions, whereas mutation of residue S1445 resulted in partial reduction of toxicity (Fig. 5A, B). To ensure that the observed loss of toxicity was not due to compromised protein stability or expression, an N-terminal 3xFLAG tag was incorporated in each of the RhsF-CT variants to allow visualisation of protein levels via immunoblotting analysis. Wild type FLAG-RhsF-CT was only detected upon co-expression with the immunity protein RhsFi to prevent toxicity. Under this condition, the levels of all RhsF-CT variants were similar (Fig. 5C). In the absence of the immunity protein, non-toxic variants R1403A and Y1456A remained present at similar levels, whereas a reduction in E1497A levels was observed, consistent with its slight residual toxicity. Taken together, these results suggest that RhsF-CT is likely an ADP-ribosyltransferase with a catalytic triad composed of residues R-Y-E.

**Figure 5.**
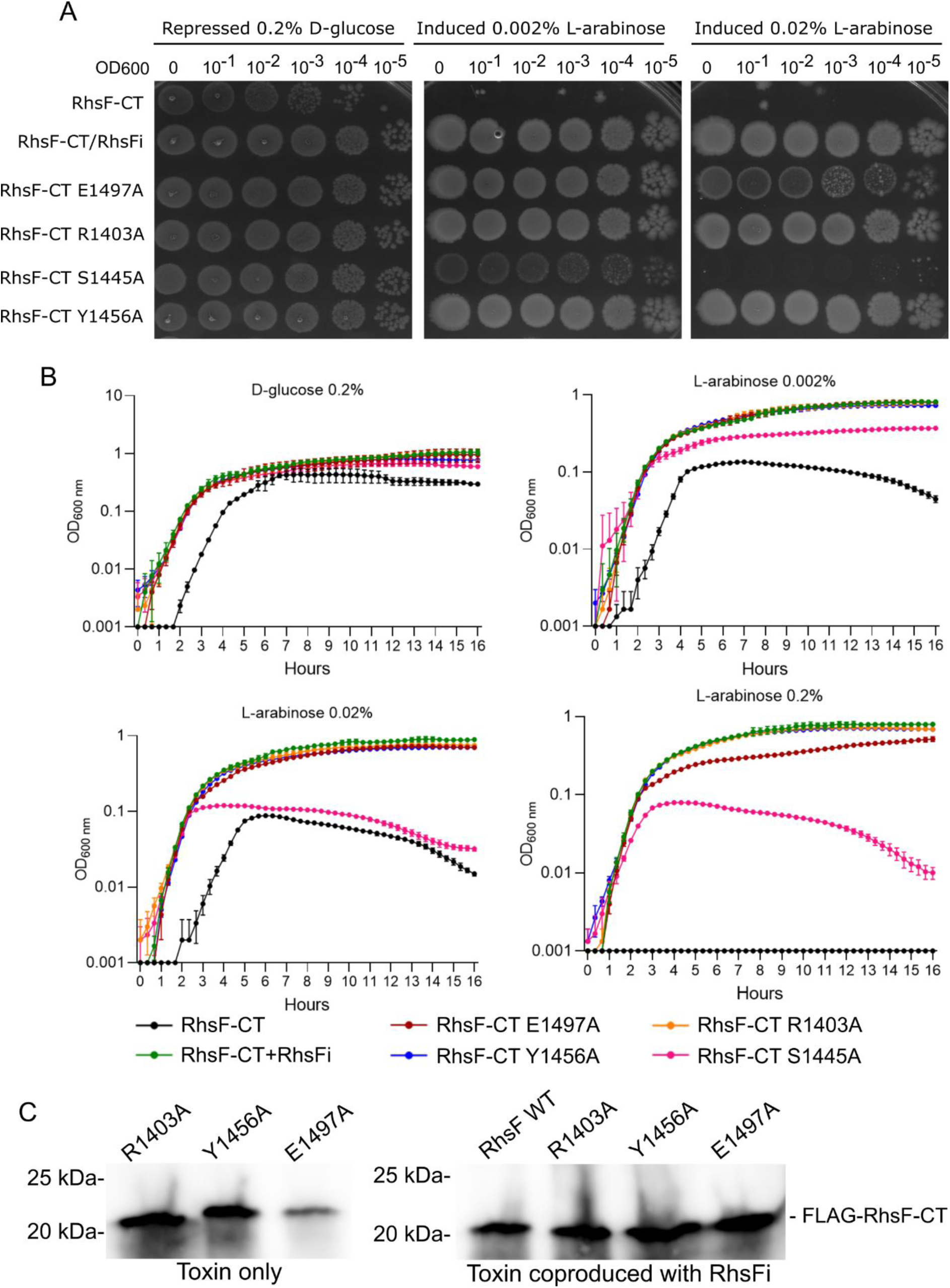
Substitution of amino acids predicted to be important for enzymatic function of RhsF-CT results in a loss of toxicity. *A-B,* growth of *E. coli* MG1655 carrying pBAD18-Kan-based plasmids directing the expression of wild type RhsF-CT alone or co-expressed with RhsFi, or of RhsF-CT variants with the amino acid substitutions indicated on solid (A) or in liquid (B) LB media containing D-glucose or L-arabinose for repression or induction, respectively, of gene expression. *C,* total protein samples from *E. coli* carrying plasmids directing the expression of RhsF-CT variants with an N-terminal 3xFLAG epitope tag (FLAG-RhsF-CT), with or without the immunity protein RhsFi, were subjected to immunoblotting analysis to visualise levels of RhsF-CT in each case.

## Discussion

In this study, we performed a global analysis of effectors secreted by the T6SS of *C. violaceum*. Combining two complementary mass spectrometry approaches, we identified six novel effector candidates, namely four phospholipases (CV_0012, CV_1234, CV_3971, and CV_3990), a hypothetical protein (CV_2125), and an uncharacterized Rhs protein (CV_1431). These findings reveal the broad repertoire of T6SS effectors employed by *C. violaceum* for interbacterial competition. Among these T6SS-associated effectors, we focused on an Rhs toxin-immunity pair, RhsF-RhsFi. We demonstrate that the FIX-containing RhsF protein is a potent T6SS-dependent antibacterial toxin that exhibits structural and catalytic features of ADP-ribosyltransferases and whose toxic activity is blocked by the cognate immunity protein RhsFi via direct occlusion of its catalytic site.

Our combined proteomics analysis was effective in identifying: (i) effectors predicted previously *in silico* (Tle1 and Tle5 phospholipases and the Rhs protein CV_1431), which are encoded within the main T6SS cluster and the four orphan VgrG clusters (Russell et al., 2013; Alves et al., 2022); (ii) a new potential cell-wall acting effector (CV_2125); (iii) Tap-1/TEC adaptor proteins associated with effector secretion (Unterweger et al., 2015; Liang et al., 2025); (iv) two PAAR proteins not identified previously (CV_2124 and CV_2664), which are encoded outside of the *C. violaceum* T6SS clusters; and (v) all six *C. violaceum* VgrG proteins (Fig. 1, Tables 1, 2). The CV_2124-CV_2126 and CV_2664-CV_2666 gene clusters likely form two operons encoding PAAR proteins (CV_2124 and CV_2664), putative cell-wall-acting effectors (CV_2125, which lacks predicted domains, and CV_2665, which contains a soluble lytic transglycosylase domain), and their putative signal peptide-containing immunity proteins (CV_2126 and CV_2666). We are currently investigating the function of these gene clusters. In a previous study, we showed that deletion of VgrG3, among all six VgrGs, has the greatest impact on T6SS firing and T6SS-mediated bacterial killing (Alves et al., 2022), but it was uncertain whether all six VgrGs were expressed and functional. Here, our data provide evidence that all six VgrG proteins are either secreted and/or interact with VgrG3 (Tables 1, 2), confirming that *C. violaceum* deploys all available VgrGs through its single T6SS. Since many cargo effectors are dependent on or show preference for specific VgrGs (Hachani et al., 2014; Cianfanelli et al., 2016), encoding six VgrGs provides *C. violaceum* with a strategy to deliver a broad range of cargo effector toxins.

We provide evidence that RhsF is a toxin delivered by the T6SS that *C. violaceum* employs for interbacterial competition (Fig. 2). Rhs proteins secreted via the T6SS are typically specialized effectors that contain a PAAR domain at their N-terminus and/or are encoded alongside chaperones that mediate interaction with VgrG proteins or less commonly are fused to VgrGs (Alcoforado Diniz and Coulthurst, 2015; Cianfanelli et al., 2016; Pei et al., 2020; Jurėnas et al., 2021; Günther et al., 2022; Kielkopf et al., 2024). In contrast, we did not identify any chaperone genes adjacent to or within the *rhsF* operon, nor detect any candidate RhsF-associated chaperones in our mass spectrometry. Furthermore, the N-terminal region of RhsF lacks a PAAR domain, and has instead a FIX (Found in type sIX) domain (Fig. 2). The FIX domain is found in the N-terminal or central regions of several predicted T6SS-secreted effectors and has been shown to be essential for the secretion, but not the toxicity, of a T6SS-dependent DNase effector from *V. parahaemolyticus* (Jana et al., 2019). Here, we show that an Rhs protein containing a FIX domain is secreted in a T6SS-dependent manner, providing experimental evidence that FIX can replace PAAR to direct secretion of Rhs-containing effectors by the T6SS. Our co-immunoprecipitation data suggests a direct interaction between RhsF and VgrG3, which contains a transthyretin-like (TTR-like) domain at its C-terminus. The TTR-like domain or an Ig-like variant domain found in many VgrG proteins have been implicated in specific interactions with the phospholipase Tle1 (Flaugnatti et al., 2016) and the Rhs effector Tse15 (Hayes et al., 2024), allowing their secretion. These findings suggest the possibility of a conserved effector-VgrG interaction mechanism mediated by TTR-like or structurally related domains. Further investigations are required to determine whether the secretion of RhsF via the T6SS machinery involves direct interaction of its N-terminal FIX domain with the TTR-like domain of VgrG3.

Our atomic structure of the C-terminal toxic domain (RhsF-CT) of RhsF in complex with the immunity protein RhsFi revealed that RhsF-CT shares strong structural similarity with ADP-ribosyltransferases (ARTs) and provided structural insights on the mechanism of toxin inhibition via direct occlusion of its catalytic site by the antitoxin RhsFi (Figs. 3, 4). Toxicity assays using point-mutated versions of the RhsF-CT underscored the importance of three key residues: R1403, Y1456 and E1497 (Fig. 5). Bacterial ARTs are classified into two classes: class I, whose catalytic site consists of the H-Y-E triad, and class II, characterized by an R-S-E triad. In both cases, there is an ultra-conserved glutamate residue (E) essential for nearly all bacterial ARTs, and a nucleophilic histidine or arginine, typically preceded by a tyrosine residue (Domenighini et al., 1994; Simon et al., 2014). Therefore, the RhsF-CT shares characteristics with ARTs of class I, but with an arginine instead of the classical histidine. Although ARTs are traditionally classified into these two main groups, bacterial ARTs with slightly divergent catalytic motifs have been reported (Baysarowich et al., 2008; Simon et al., 2014; Bullen et al., 2022), including the type III secretion system (T3SS) effector CteC, an ART of *C. violaceum* that ADP-ribosylates host ubiquitin during infection (Yan et al., 2020; Tang et al., 2024). The C-terminal toxic domains of Rhs proteins are very variable, including DNases, NADases, and ADP-ribosyltransferases (Koskiniemi et al., 2013; Alcoforado Diniz and Coulthurst, 2015; Jurėnas et al., 2021; Bullen et al., 2022; Jurėnas et al., 2022; Hagan et al., 2023). While RhsF-CT displays catalytic and structural features of an ART toxin, further studies are required to demonstrate its ADP-ribosyltransferase activity on a molecular target which triggers death or stasis during interbacterial competition. Overall, we have characterised an important component of the antibacterial effector repertoire of the *C. violaceum* T6SS and demonstrated T6SS-mediated deployment of a FIX-containing Rhs protein. Our findings contribute to a deeper understanding of how bacteria utilise T6SSs and diverse antibacterial effectors to compete against rival bacteria and shape polymicrobial communities.

## Experimental procedures

### Bacterial strains, plasmids, and growth conditions

Bacterial strains and plasmids used in this study are listed in Table 4 and Table S2, respectively. *C. violaceum* and *E. coli* strains were cultured at 37 °C in Luria-Bertani (LB) (10 g/L tryptone, 5 g/L yeast extract, 10 g/L NaCl) liquid medium with shaking at 200 rpm or on solid LB medium (supplemented with 15 g/L agar). When necessary, antibiotics were added at the following concentrations: ampicillin 100 μg/ml, kanamycin 50 μg/ml, nalidixic acid 20 μg/ml, chloramphenicol 25 μg/ml, and streptomycin 100 μg/ml. To repress gene expression from pBAD18-Kn, 0.5% D-glucose was added to the media for cloning and maintenance. The minimal medium used for proteomic analysis of *C. violaceum* supernatant had the following composition: phosphate buffer (K₂HPO₄ 40 mM, KH₂PO₄ 8 mM), ammonium sulfate 0.1%, magnesium sulfate 0.41 mM, glucose 0.4%, Complete Supplement Mixture (CSM) Single Drop-Out: -Adenine (Formedium) 780 mg/l, and adenine 0.01%. Solid minimal medium was prepared by supplementing with 15 g/L agar. DNA cloning procedures in plasmids were performed using *E. coli* DH5α, except for constructs with the pKNG101 vector, which were generated using *E. coli* CC118λpir. The transfer of plasmid constructs into *C. violaceum* strains was carried out via triparental conjugation (*E. coli* CC118λpir along with *E. coli* HH26 (pNJ5000) as the helper strain).

**Table 4.**
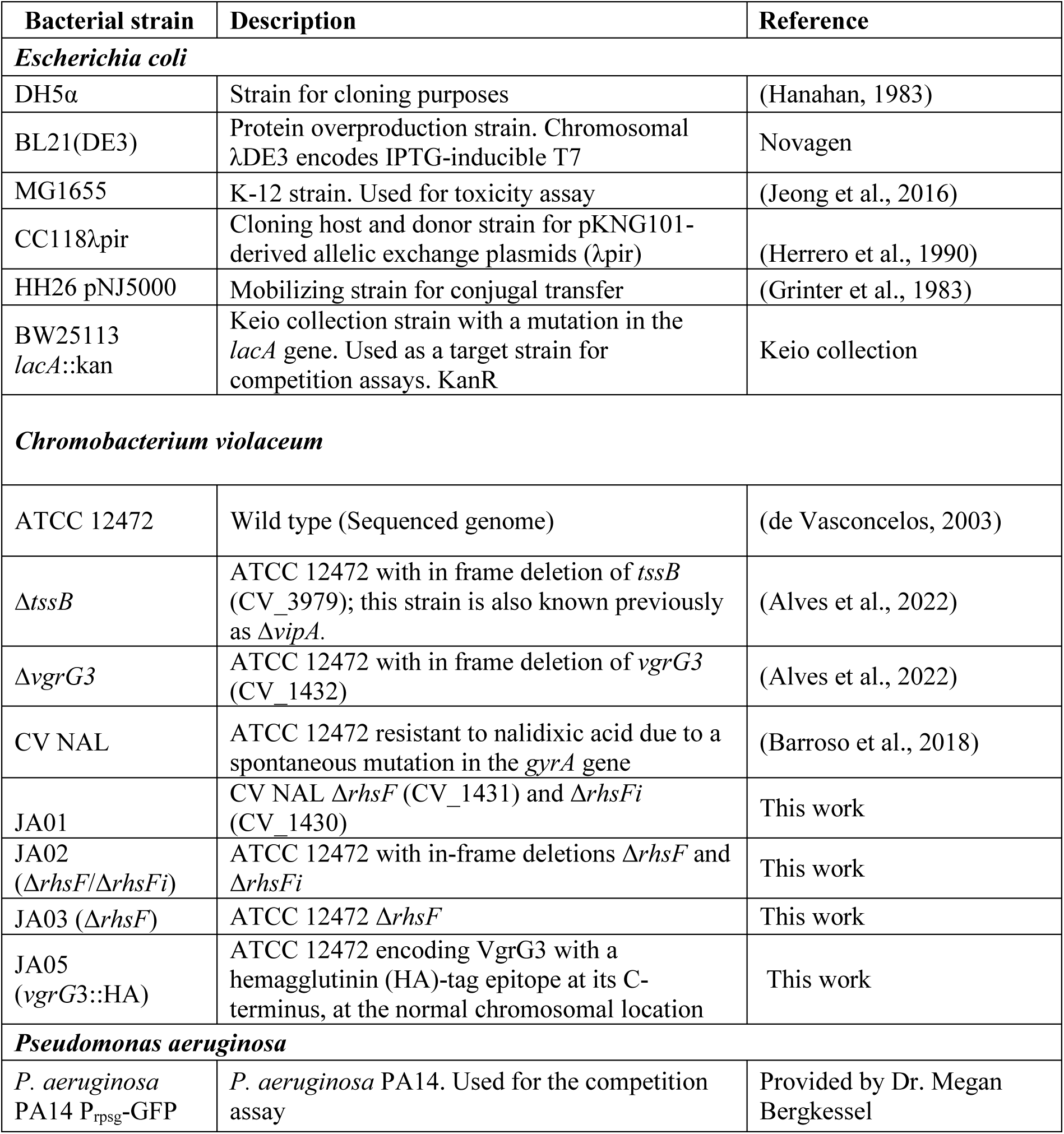
Bacterial strains used in this work.

### Construction of C. violaceum mutant strains

Null mutant strains of *C. violaceum* (non-polar in-frame deletion of a given gene) were generated by allelic exchange through homologous recombination (Barroso et al., 2018; Batista et al., 2020; Alves et al., 2022) using the suicide vector pKNG101 (Kaniga et al., 1991; Murdoch et al., 2011). The flanking regions of the target gene were amplified by PCR using specific primers (Table S3), joined following Gibson Assembly® Master Mix Protocol, and cloned into pKNG101 vector. The resulting constructs were transferred to *C. violaceum* by triparental conjugation. Transconjugants were selected on LB agar plates containing ampicillin and streptomycin (for pKNG101) to select the first homologous recombination event. Isolated colonies were grown in LB and plated on LB supplemented with 16% sucrose for counter-selection of the second recombination event. Deletion of the target genes was verified by PCR and DNA sequencing. For the construction of strain used in the co-immunoprecipitation assay (JA05), a hemagglutinin (HA) tag was inserted upstream of the *vgrG*3 stop codon in the *C. violaceum* genome by homologous recombination via plasmid pSC3913, and confirmed by PCR.

### Interspecies interbacterial competition

The interbacterial competition assay was performed as previously described (Murdoch et al., 2011; Alves et al., 2022). Attacker (*C. violaceum*) and target strains were grown overnight at 37 °C on LB agar plates, resuspended in liquid LB, normalized to an OD600 of 5 and mixed at a 5:1 ratio (attacker:target). Then, 10 μl of the mixture were spotted onto LB agar plates and incubated for 4 hours at 37 °C to allow competition. After this period, cells were recovered in 1 ml of LB, serially diluted, and plated on LB agar containing kanamycin (to select *E. coli* BW25113 *lacA*::kan) or nalidixic acid (to select *P. aeruginosa* PA14) to determine the CFU (colony-forming units) of the target strains.

### Intraspecies bacterial competition

Intraspecies competition assays were performed to assess susceptibility to T6SS effector-mediated attack between sibling *C. violaceum* cells. For this, nalidixic acid-sensitive *C. violaceum* strains (attacker) and nalidixic acid-resistant strains (target) were used. Attacker and target strains were grown overnight at 37 °C on LB agar plates, resuspended in liquid LB, normalized to an OD600 of 0.5, and mixed at a 1:1 ratio. Then, 10 μl of the mixture were spotted onto LB agar plates, which were incubated at 37 °C for 7 hours to allow competition. After this period, cells were recovered in 1 ml of LB, serially diluted, and plated on LB agar containing nalidixic acid to determine the CFU (colony-forming units) of the target strains.

### Toxicity assay assessed by spot viability, growth curves, and light microscopy

The pBAD18-Kn expression vector, either empty or containing inserts to allow the expression of toxins with or without their corresponding immunity proteins, was used to transform *E. coli* MG1655. Transformants were grown overnight at 37 °C in LB liquid medium supplemented with kanamycin and 0.5% D-glucose to repress expression of the cloned genes. Cultures were normalized to an OD600 of 1, from which 10-fold serial dilutions were prepared in PBS (phosphate buffered saline). For spot viability, 5 μl of each dilution were spotted onto LB agar plates containing kanamycin and either 0.2% D-glucose or L-arabinose at concentrations of 0.002% or 0.02%. After drying, plates were incubated at 37 °C overnight before image acquisition.

For growth curves, overnight cultures grown in LB medium supplemented with kanamycin and 0.5% D-glucose were used to inoculate cultures to a starting OD600 of 0.01 in LB medium containing kanamycin and either 0.5% D-glucose to repress gene expression or L-arabinose at 0.002%, 0.02%, or 0.2% to induce expression. Growth was monitored in a 96-well plate using a BioTek Synergy HTX Multimode Reader at 37 °C with vigorous and continuous shaking.

For time-lapse microscopy, *E. coli* MG1655 strains carrying pBAD18-Kn-based constructs directing the expression of *rhsF*-CT or *rhsF*-CT/*rhsFi* were grown overnight in LB medium with kanamycin and 0.5% D-glucose, and then these cultures were diluted in liquid LB to an OD600 of 0.05. A 10 μl aliquot was pipetted onto a coverslip coated with solid LB medium supplemented with kanamycin and 0.02% L-arabinose. Images were captured for up to 8 hours at 3- to 5-minute intervals using a Nikon Biostation IM-Q light microscope, which maintains the samples at 37 °C under controlled humidity conditions.

### Immunoblotting to determine stability of RhsF-CT variants and expression of VgrG3-HA

Genes encoding variants of RhsF-CT with an N-terminal 3xFLAG epitope tag, with or without RhsFi, were cloned in pBAD18-Kan (Tables S2, S3). Overnight cultures grown in LB media supplemented with kanamycin and D-glucose 0.5% were used to inoculate cultures to a starting OD600 of 0.02 in LB media supplemented with kanamycin. Cells were grown for 2 hours at 37 °C, then L-arabinose was added to a final concentration of 0.2% for a further 2 hours growth. Then, 1 ml of cultures were harvested by centrifugation, combined with 4X protein loading buffer (Tris-HCl 200 mM pH 6.8, EDTA 6.4 mM, glycerol 32%, SDS 6.4% and bromophenol blue 0.007%) at a volume proportional to the culture OD600 (100 µl of buffer for OD600=1) and heated at 100 °C for 10 minutes. For detection of the VgrG3-HA, protein samples obtained following the co-immunoprecipitation protocol (see section below) were mixed at a 1:1 volume ratio with 4X protein loading buffer. Proteins were separated by 4-20% SDS-PAGE and transferred onto PVDF membrane (Millipore) via dry transfer using the iBlot2 system (Thermo Fisher). The membrane was blocked with milk powder 2.5% (Marvel) suspended in PBS + 0.1% Tween-20. Membrane washing was done with PBS + 0.1% Tween-20. 3xFLAG-RhsF-CT was detected using the primary antibody, anti-FLAG (Sigma Aldrich, #F3165) at 1:10000, and the VgrG3-HA using an anti-HA monoclonal antibody (Sigma Aldrich, #H6533) at 1:5000. The secondary antibody, HRP-conjugated anti-mouse (BioRad, #170-6516) was used at 1:10000. For signal detection, the membrane was treated with HRP substrate (Millipore) and imaged using an Azure 600 imaging system.

### Preparation of samples for analysis of the C. violaceum supernatant by mass spectrometry

To identify T6SS-secreted effectors, *C. violaceum* wild type and Δ*tssB* strains were grown in seven biological replicates overnight in LB at 37 °C. Cultures were diluted to an OD600 of 0.04 in 250 ml of minimal medium, and incubated for 8 hours until reaching OD600 of 1.0. Cultures were subjected to centrifugation at 5000 xg for 30 minutes at 4 °C, and the culture supernatant was carefully collected for another round of centrifugation. This process was repeated four times to minimize cellular contamination. After the fourth centrifugation, 100 ml of culture supernatant was collected and treated with 6.25% TCA (trichloroacetic acid) for 14 hours at 4 °C. Precipitated proteins were recovered by centrifugation at 5000 xg for 30 minutes at 4 °C, and the resulting pellet was washed five times with 80% cold acetone. Protein samples were then dried under laminar airflow for 30 minutes.

### Proteomics sample preparation, LC-MS, analysis and post-processing

Precipitated protein samples were resuspended in 50 mM triethyl ammonium bicarbonate (TEAB) pH 8.5 with 5 % (w/v) SDS and 1x complete protease inhibitor cocktail (Roche). Samples were sonicated using a UP200St ultrasonic processor (Hielscher) at 90 W, 45 s pulse, 15 s rest, three times. Protein concentration was quantified using Pierce BCA Protein Assay (Thermo Fisher). Samples (30 µg) were then denatured with 5 mM TCEP at 60 °C for 15 minutes, alkylated with 30 mM iodoacetamide at room temperature (20 °C) for 30 minutes in the dark, and acidified to a final concentration of 2.5 % phosphoric acid. Samples were then diluted eightfold with 90 % MeOH 10 % TEAB (pH 7.2) and added to the S-trap (Protifi) micro columns. The manufacturer-provided protocol was then followed, with a total of five washes in 90 % MeOH 10 % TEAB (pH 7.2), and trypsin added at a ratio of 1:10 enzyme:protein and digestion performed for 2 h at 47 °C. After elution, peptides were dried using a vacuum concentrator and stored at -80 °C.

Immediately before mass spectrometry, samples were resuspended in 2% acetonitrile 0.1% trifluoroacetic acid in LC-MS grade H2O, and each sample was independently analysed on an Orbitrap Exploris 480 mass spectrometer (E480, Thermo Fisher), connected to an UltiMate 3000 RSLCnano System (Thermo Fisher). Peptides (1 µg) were injected on a PepMap 100 C18 LC trap column (300 μm ID × 5 mm, 5 μm, 100 Å) followed by separation on an EASY-Spray nanoLC C18 column (75 μm ID × 50 cm, 2 μm, 100 Å). Solvent A was water containing 0.1% formic acid, and solvent B was 80% acetonitrile containing 0.1% formic acid. The gradient used for analysis of proteome samples was as follows: solvent B was increased from 3-5 % for 5 min, then from 5-35 % for 60 mins, then increased to 90 % for 2.5 mins, held at 95 % for 2.5 mins, then decreased to 3 % B in 0.5 mins and held at 3 % B for 5 mins. Flow was kept constant at 400 nl/min. The E480 was operated in positive mode DIA MS2. A precursor ion scan (full scan) was performed in the range 350-1050 m/z, 60000 resolution, normalised AGC target of 150 %, RF lens set to 50 %. MS2 scans were performed at 30000 resolution, with isolation windows of 2 m/z, HCD normalised collision energy set to 30 %, time to 50 ms. Overall cycletime was 3 seconds.

Raw files were searched using DIA-NN V 1.8 (Demichev et al., 2020), using its *in silico* generated spectral library function, based on reference proteome FASTA files for *C. violaceum* (UP000001424, downloaded from UniProt on 21/07/2023) and a common contaminants list (Frankenfield et al., 2022). Trypsin specificity with a maximum of 1 missed cleavage was permitted per peptide, cysteine carbamidomethylation was set as a fixed modification, methionine oxidation as variable, maximum variable modifications was set to 1. All other variables were left as defaults-protein and peptide false discovery rate (FDR) were set to 1 %. Raw data and DIA-NN results files were uploaded to the proteomeXchange via the PRIDE partner repository, under PXD071363. Post-processing was performed in R: The DIA-NN output pg.matrix was processed to include only proteins identified by at least 2 peptides, contaminants were auto-excluded. Protein log2 intensity values were subjected to processing with limma (Ritchie et al., 2015), with significance inferred by FDR corrected T-test (P = <0.05) and fold change of ±2-fold.

### Immunoprecipitation of VgrG3-HA and mass spectrometry analysis of associated proteins

Co-immunoprecipitation followed by mass spectrometry (Co-IP/MS), was conducted in biological triplicates. Overnight cultures of wild type *C. violaceum* (no HA tag, negative control) and *vgrG*3-HA strains were used to inoculate cultures to a starting an OD600 of 0.05 in 50 ml of LB medium and grown at 37 °C until reaching an OD600 of 2.5. Cells were harvested by centrifugation at 48000 xg for 20 minutes at 4 °C, and resuspended in 5 ml of lysis buffer (20 mM Tris-Cl pH 7.5, 150 mM NaCl, 0.5 mM EDTA, 0.1% Triton X-100, and protease inhibitor cocktail (Roche). Cell lysis was performed by sonication, and the soluble fraction was obtained by centrifugation at 21000 xg for 20 minutes at 4 °C. For Co-IP of proteins using magnetic beads, 30 μl of HA-tag Rabbit mAb magnetic beads (NEB) were washed three times with wash buffer (20 mM Tris-Cl pH 7.5, 150 mM NaCl, 0.5 mM EDTA, 0.1% Triton X-100), and incubated with the entire soluble fraction for 16 hours at 4 °C under gentle rotation. After incubation, the supernatant was discarded, and the beads were washed three times. Protein elution was performed using 30 μl of elution buffer (5% SDS, 50 mM Tris pH 7.5, 150 mM NaCl) incubated at 70 °C for 5 minutes.

Sample preparation and trypsin digestion for mass spectrometry were performed at the Proteomics Facility of the Centre for Advanced Scientific Technologies, University of Dundee. Protein concentrations were determined using the Micro-BCA assay (Thermo Scientific) and were processed using S-Trap Micro columns (Protifi) where proteins were reduced, alkylated and digested overnight at 37°C at 1:40 enzyme-to-substrate. A second digest was repeated for 6 hours the following day. Digested peptides were run on a Q-Exactive Plus (Thermo Scientific) instrument coupled to a Dionex Ultimate 3000 HPLC system (Thermo Scientific) with LC buffers compromising of buffer A (0.1% formic acid) and buffer B (80% acetonitrile, 0.1% formic acid). The buffers were used to create a gradient lasting 185 minutes where the peptides were eluted from a 200 cm uPAC nanoLC C18 column (Pharma Fluidics) at a flow rate of 300 nl/min for the first 156 minutes before increasing to 400 nl/min for the remainder of the gradient. Raw data was acquired in Data Independent Acquisition (DIA) mode. A scan cycle compromised a full MS scan with an m/z range of 345-1155, resolution of 70,000, Automatic Gain Control (AGC) target 3x10^6^ and a maximum injection time of 200 ms. MS scans were followed by DIA scans of dynamic window widths with an overlap of 0.5 Th. DIA spectra were recorded with a first fixed mass of 200 m/z, resolution of 17,500, AGC target 3x10^6^ and a maximum IT of 55 ms. Normalised collision energy was set to 25 % with a default charge state set at 3. Data for both MS scan and MS/MS DIA scan events were acquired in profile mode. Peptides were initially trapped on an Acclaim PepMap C18, (100µm x 2cm) and then separated on a 200 cm uPAC nanoLC C18 column (Pharma Fluidics). The column was kept at a constant temperature of 50 °C and a source voltage of 2.0 kV. Data are available via ProteomeXchange with identifier PXD071369. Label-free analysis was performed in MaxQuant version 1.6.2.10 using the RAW files. Further data analysis was performed using Perseus version 1.6.12.0 to generate p-values and fold change and perform student t-tests.

### Cloning, expression, and purification of the RhsF-CT/RhsFi complex

The RhsF-CT/RhsFi complex was expressed using the pACYCDuet-1 expression vector, which contains two multiple cloning sites (MCSs) and is designed for the co-expression of two genes. The DNA fragment encoding the full-length RhsFi immunity protein was cloned into MCS-2. Subsequently, the DNA fragment encoding the toxic C-terminal portion of RhsF (RhsF-CT, starting from amino acid 1393 of RhsF) plus His6Tag and TEV protease recognition sequence was amplified from pET15b and cloned into MCS-1. For protein production, *E. coli* BL21(DE3) was freshly transformed with pSC3912 and the transformed cells were grown in liquid LB medium supplemented with chloramphenicol at 30 °C until reaching an OD600 of 0.8. The temperature was then reduced to 20 °C, and expression was induced overnight with 1 mM isopropyl β-D-1-thiogalactopyranoside (IPTG). Cells were harvested by centrifugation and resuspended in lysis buffer (50 mM Tris-HCl pH 8.0, 250 mM NaCl, 30 mM imidazole, and 0.5 mM TCEP), supplemented with DNaseI (Sigma Aldrich) and an EDTA-free protease inhibitor cocktail (Sigma Aldrich). Cell lysis was performed using a cell disruptor at 30 pounds per square inch (PSI). The soluble fraction was obtained by centrifugation at 40.000g for 30 minutes at 4 °C and clarified with 0.22 µm filters.

The His6-RhsF-CT/RhsFi complex was purified using a 5 ml Cytiva Ni^2+^ HisTrap HP column (GE healthcare). After loading the soluble fraction, the column was washed with lysis buffer, and the complex was eluted using linear gradient of imidazole. Desalting was then performed using a HiPrep 23/10 Desalting column into storage buffer (50 mM Tris-HCl pH 8.0, 250 mM NaCl, 0.5 mM TCEP). For X-ray crystallography, the His6-tag was removed by treating purified complex with TEV protease (1 mg of TEV per 10 mg of protein) overnight at 4 °C. To eliminate any remaining His6-tagged protein, cleaved tag and TEV-protease, the TEV-digested sample was subjected to reverse purification on a HisTrap HP column (GE healthcare). Finally, the purified complex was subjected to size-exclusion chromatography (SEC) on Superdex 75 HiLoad 16/600 column (GE Healthcare) equilibrated with buffer (20 mM Tris-HCl pH 8.0, 150 mM NaCl, and 0.5 mM TCEP-HCl). The proteins were loaded with either 1.5 ml or 2 ml loops and eluted with the same buffer.

### X-ray crystallography and diffraction data collection

Crystallization condition screening for the RhsF-CT/RhsFi complex was performed using commercial 96-well screening plates from Molecular Dimensions Ltd and Qiagen. The sitting-drop vapor diffusion method was used, combining 0.1 μl of protein solution (25 mg/ml) with 0.1 or 0.2 μl of reservoir solution. Crystallization plates were incubated at 20 °C, and crystal formation was monitored under a polarized light stereomicroscope. The four crystals selected for X-ray diffraction were obtained from the PACT Premier screen (Molecular Dimensions), after supplementing the RhsF-CT/RhsFi purification buffer with 20% ethanol. These crystals were obtained under the following specific condition: Complex concentration: 8.33 mg/ml, pH 6.0; Ethanol 6.7%; PEG1500 16.7%; Tris-Cl 16.7 mM; NaCl 83.3 mM; and MMT buffer (malic acid, MES, and Tris) 67 mM.

X-ray diffraction data were collected and processed at the Diamond Light Source (DLS) synchrotron facility in Didcot, UK, using standard protocols. Reservoir solutions supplemented with 20–25% (v/v) glycerol were used as cryoprotectants for the four selected RhsF-CT/RhsFi crystals. Individual crystals were frozen and diffraction data was collected at beamline I03 using an Eiger2 XE 16M detector. Crystals were maintained at 100 K using a gaseous nitrogen stream. For each crystal, 360 diffraction images were collected with an oscillation of 0.1° per image. The crystal-to-detector distances were 200 mm (for crystals 1 and 3) and 290 mm (for crystals 2 and 4), using an X-ray wavelength of 0.9763 Å. The collected diffraction data were indexed, integrated, and scaled using XDSgui (Brehm et al., 2023). The Matthews coefficients (Å3.Da-1) and solvent content (%) were calculated using Xtriage within the Phenix software suite (Liebschner et al., 2019).

For structure refinement, the top-ranked ColabFold (Mirdita et al., 2022) model of the RhsF-CT/RhsFi complex was iteratively refined and manually adjusted using Phenix (Liebschner et al., 2019) and Coot (Emsley et al., 2004), respectively. After initial refinement cycles, water molecules were added automatically by Phenix and manually verified based on electron density shape and hydrogen bonding potential. Throughout refinement, model geometry, structural quality, and fit to experimental data were monitored in Coot and evaluated using key parameters such as Rfree, Rwork, clashscore, RMSD (bond lengths and angles), and B-factors via MolProbity (Chen et al., 2010) and other Phenix tools (e.g., POLYGON) (Urzhumtseva et al., 2009). This refinement procedure was performed separately for all four datasets, with progressive reduction in Rfree values. The dataset yielding the highest data completeness and lowest values for resolution, Rfree, Rwork, and clashscore, alongside angle RMSD close to 1 and bond RMSD near 0.01, was selected as the final model. Structural figures were generated using Chimera and ChimeraX (Meng et al., 2023).

## Supporting information

four Supporting Figures and three Supporting Tables

## Data availability

The crystal structure of the RhsF-CT/RhsFi complex has been deposited into the Protein Data Bank (PDB) under the accession code 9ZDM. The mass spectrometry proteomics data are available in the ProteomeXchange Consortium via the PRIDE partner repository with the dataset identifier PXD071363 for the secretome analysis and PXD071369 for the co-immunoprecipitation of VgrG3. All other data are included within the manuscript or the supporting information.

## Supporting information

This article contains supporting information (four Supporting Figures and three Supporting Tables).

## Acknowledgments

We would like to acknowledge the Diamond Synchrotron Light source for beamtime and the expert assistance from the FingerPrints Proteomics Facility, University of Dundee. We thank Dr. Megan Bergkessel for providing the *Pseudomonas aeruginosa* PA14 Prpsg-GFP strain and Roberta Rosales for technical support in microscopy facility. For the purpose of Open Access, the authors have applied a CC BY public copyright licence to any Author Accepted Manuscript version arising from this submission.

## Author contributions

J.A.A and G.P. performed the experimental work. J.A.A., G.P., S.J.C., and J.F.S.N. performed experimental design, data analysis and interpretation. A.M.F. and M.T. performed the mass spectrometry analysis of supernatants. G.G.S. refined and deposited the structure of RhsF/RhsFi complex. J.A.A. and J.F.S.N. wrote the manuscript with contributions and input from the other authors. J.F.S.N. and S.J.C. participate in study coordination and funding acquisition. All authors read and approved the final manuscript.

## Funding and additional information

This study was financed, in part, by the São Paulo Research Foundation (FAPESP; process numbers 2020/00259-8, 2021/06894-0, and 2021/10577-0), Brasil, and Fundação de Apoio ao Ensino, Pesquisa e Assistência do Hospital das Clínicas da FMRP-USP (FAEPA). During this work, J.A.A. was supported by fellowships from FAPESP (2018/03979-1 and 2022/01586-8).

J.F.S.N. is Research Fellow from CNPq (Conselho Nacional de Desenvolvimento Científico).

This work was also supported by Wellcome (grant numbers 215599/Z/19/Z, S.J.C, 220321/Z/20/Z, S.J.C.).

## Abbreviations

ADP: adenosine diphosphate
His6: hexahistidine tag
HRP: horseradish peroxidase
OD600: optical density at 600 nm
TCEP: tris(2-carboxyethyl)phosphine

