## Supplementary material for "Global identification of *Chromobacterium violaceum* T6SS effectors reveals an Rhs antibacterial toxin featuring FIX and ADP-ribosyltransferase domains": four Supporting Figures and three Supporting Tables

**Figure S1. Strategies for identification of *C. violaceum* T6SS effectors.**

**Figure S2**. **Time-lapse microscopy of *E. coli* expressing RhsF or RhsF/RhsFi.**

**Figure S3**. **Crystallization of the RhsF-CT/RhsFi complex.**

**Figure S4. Residues involved in the protein interface between RhsF-CT and RhsFi.**

**Table S1.** List of proteins showing structural similarity with RhsF-CT.

**Table S2**. Plasmids used in this work.

**Table S3**. Oligonucleotide primers and synthetic gene fragments.

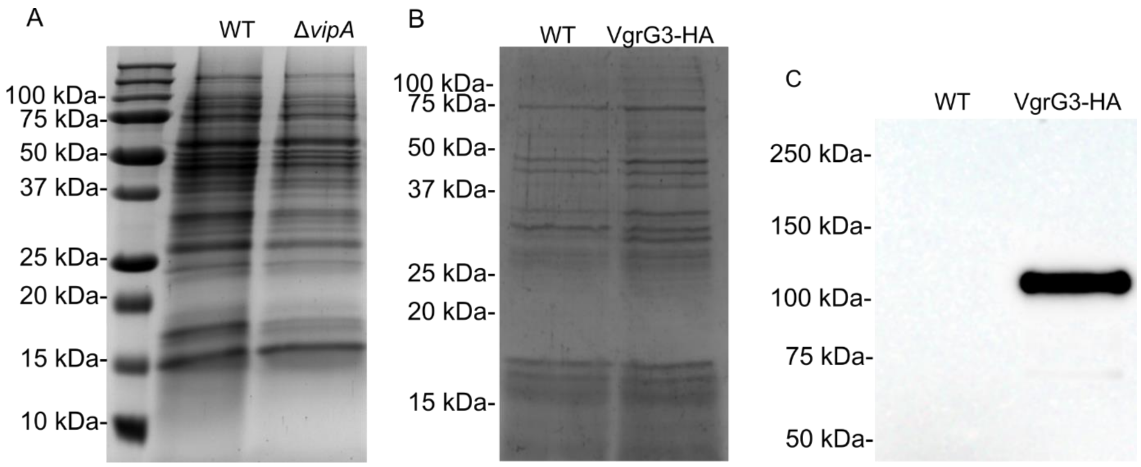


**Figure S1. Strategies for identification of *C. violaceum* T6SS effectors.** *A,* proteins secreted by *C. violaceum* wild-type (WT) and Δ*tssB* strains, obtained by TCA precipitation and resolved by SDS-PAGE. *B,* proteins eluted from an anti-HA immunoprecipitation performed on cell lysates of wild type *C. violaceum* (negative control) or the strain encoding VgrG3-HA, resolved by SDS-PAGE and stained with Coomassie Blue. *C,* anti-HA immunoblotting, confirming the expression of the VgrG3-HA fusion protein (107 kDa).

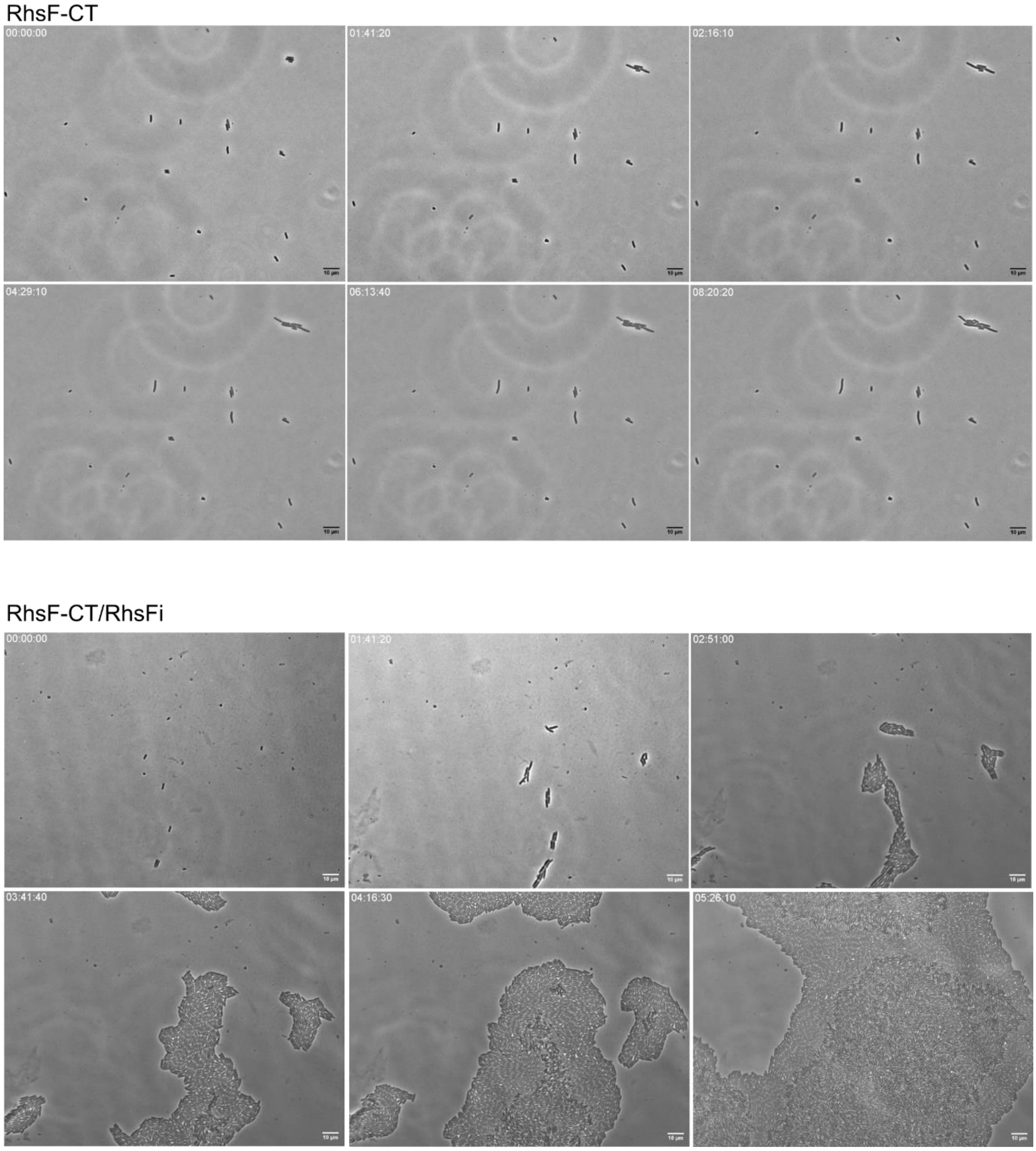


**Figure S2**. **Time-lapse microscopy of *E. coli* expressing RhsF or RhsF/RhsFi.** Light microscopy of *E. coli* MG1655 carrying the pBAD18-Kan vector expressing the toxic C-terminal domain of RhsF (RhsF-CT) or co-expressing RhsF-CT with the immunity protein RhsFi. Strains were monitored for approximately 8 hours to evaluate their growth.

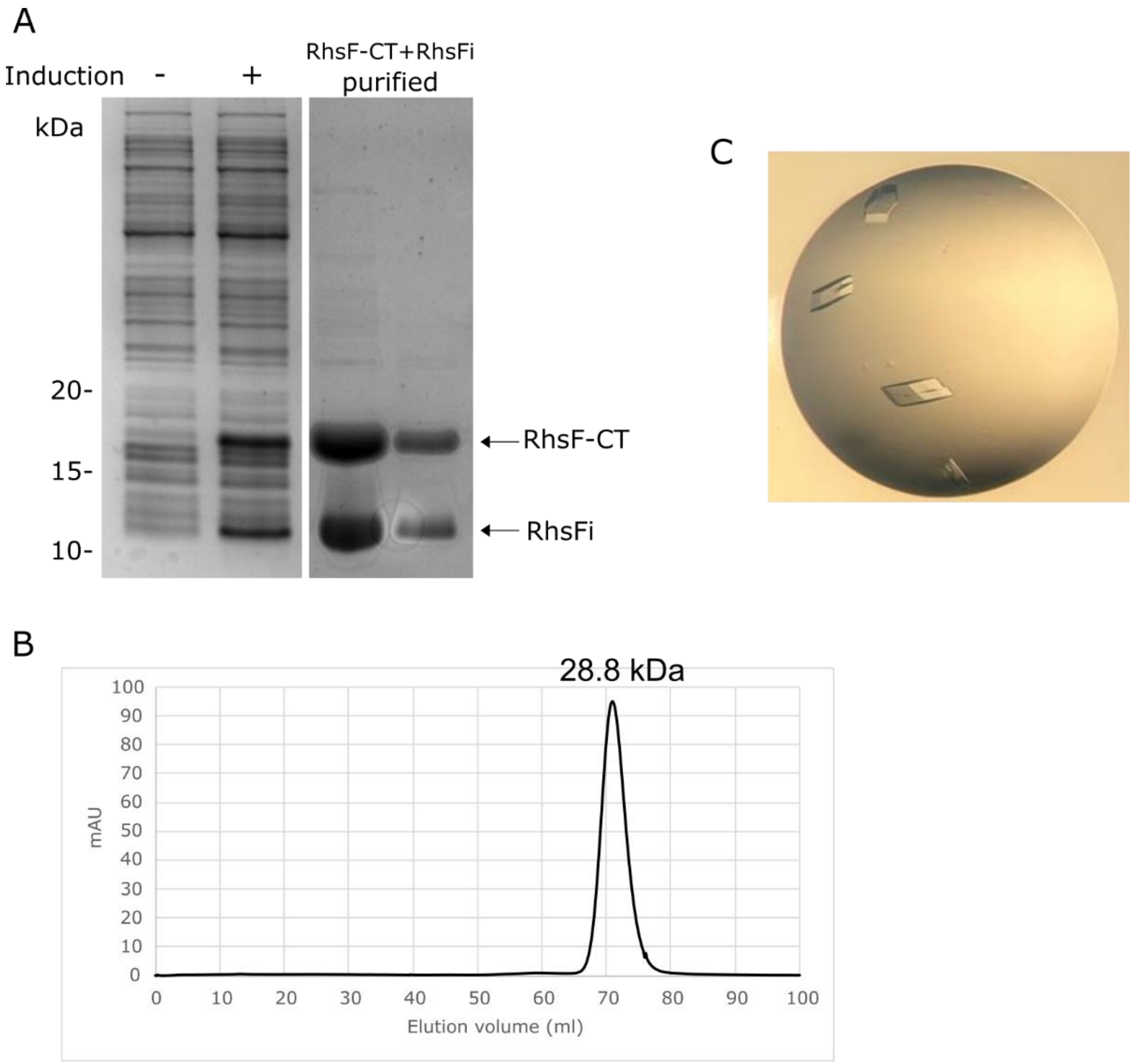


**Figure S3**. **Crystallization of the RhsF-CT/RhsFi complex.** *A,* SDS-PAGE analysis of total protein with (+) or without (-) induction of production of the RhsF-CT and RhsFi proteins by addition of IPTG to cultures of *E. coli* BL21(DE3) carrying pSC3912 (left), and of eluted proteins following nickel affinity purification of the RhsF-CT/RhsFi complex (right). *B,* size exclusion chromatography on Superdex 75 HiLoad 16/600 column (GE Healthcare) of the purified RhsF-CT/RhsFi complex. The elution volume corresponds to a complex with 28.8 kDa. *C,* image of the four RhsF-CT/RhsFi crystals obtained and submitted for X-ray diffraction analysis.

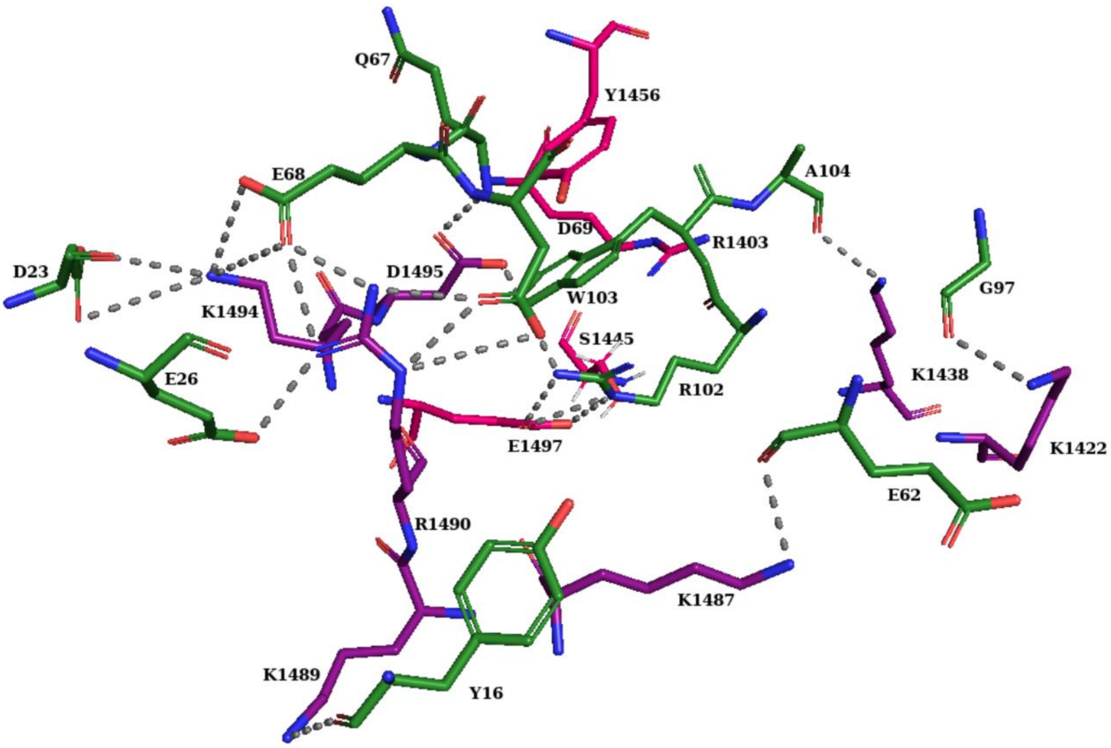


**Figure S4. Residues involved in the protein interface between RhsF-CT and RhsFi.** Carbon atoms are coloured purple for RhsF-CT and green for RhsFi, while the carbon atoms of residues E1497, S1445, Y1456, and R1403 in RhsF-CT are highlighted in pink. Salt bridges and hydrogen bonds are indicated by dashed lines. These were identified using PDBePISA (https://www.ebi.ac.uk/pdbe/pisa/) based on the crystallographic structure. Salt bridges were defined and counted per N–O atom pair with distance ≤ 4.0 Å between oppositely charged side chain.

**Table S1.** Proteins showing structural similarity with RhsF-CT.

| PDB entry | Z-score | Q-score | RMSD | Sequence identity % | Target (PDB entry) | Protein name | Organism |
|---|---|---|---|---|---|---|---|
| 5zj4:A | 7.4 | 0.34 | 1.96 | 22 | Guanine-specific ADP ribosyltransferase | ScARP | *Streptomyces coelicolor* |
| 5zj5:B | 7.3 | 0.30 | 1.98 | 23 | Guanine-specific ADP ribosyltransferase with NADH and GDP | ScARP | *Streptomyces coelicolor* |
| 4z9d:A | 8.4 | 0.28 | 2.15 | 17 | EcPltA | EcPltA | *Escherichia coli* |

Proteins identified using PDB e-Fold with RMSD (root mean square deviation) ≤ 2.5 and Z-score (standard score) ≥ 5.0 are included.

**Table S2**. Plasmids used in this work.

| Plasmids | Description | Source/Reference |
|---|---|---|
| pBAD18-Kan | Arabinose-inducible expression vector; gene of interest is cloned downstream of the Para promoter (KanR) | (Guzman et al., 2000) |
| pACYC-Duet-1 | Protein overproduction vector for the co-expression of two genes. Each multiple cloning site (MCS) is preceded by a T7 promoter, and the first site allows for fusion of an N-terminal His6 tag (KanR) | Novagen |
| pKNG101 | Suicide vector for allelic exchange (SmR, sacBR, mobRK2, oriR6K) | (Kaniga et al., 1991) |
| pET15b-TEV | Vector for protein overexpression under the control of the T7 promoter. Permits fusion of an His6 tag followed by a TEV protease cleavage site to the N-terminus of the overexpressed protein (ApR) | (Rao et al., 2011) |
| pSC3904 | pKNG101-derived allelic exchange plasmid for generation of an in-frame deletion of *rhsF* (CV_1431) and *rhsFi* (CV_1430) | This work |
| pSC3905 | pKNG101-derived allelic exchange plasmid for generation of an in-frame deletion of *rhsF* | This work |
| pSC3913 | pKNG101-derived allelic exchange plasmid for the genomic insertion of a hemagglutinin epitope at the C-terminal region of VgrG3 (CV1432) | This work |
| pSC3911 | Coding sequences for RhsF-CT and RhsFi in pET15b-TEV. RhsF fused to His6Tag and TEV sequence | This work |
| pSC3912 | Coding sequences for RhsF-CT and RhsFi in pACYC-Duet. *rhsF* fused with TEV and His6Tag was amplifyed from pSC3911 and cloned into the first MCS | This work |
| pSC3948 | Coding sequence for 3xFLAG-RhsF-CT (CV_1431; amino acids 1393- 1513) and RhsFi (CV_1430) in pBAD18-Kan. Produced by Genscript | This work |
| pSC3949 | Coding sequence for 3xFLAG-RhsF-CT R1403A and RhsFi (CV_1430) in pBAD18-Kan. Produced by Genscript | This work |
| pSC3950 | Coding sequence for 3xFLAG-RhsF-CT Y1456A and RhsFi (CV_1430) in pBAD18-Kan. Produced by Genscript | This work |
| pSC3951 | Coding sequence for 3xFLAG-RhsF-CT E1497A and RhsFi (CV_1430) in pBAD18-Kan. Produced by Genscript | This work |
| pSC3952 | Coding sequence for 3xFLAG-RhsF-CT in pBAD18-Kan. Derived from pSC3948 by digestion with *SalI* | This work |
| pSC3953 | Coding sequence for 3xFLAG-RhsF-CT R1403A in pBAD18-Kan. Derived from pSC3949 by digestion with *SalI* | This work |
| pSC3954 | Coding sequence for 3xFLAG-RhsF-CT Y1456A in pBAD18-Kan. Derived from pSC3950 by digestion with *SalI* | This work |
| pSC3955 | Coding sequence for 3xFLAG-RhsF-CT E1497A in pBAD18-Kan. Derived from pSC3951 by digestion with *SalI* | This work |

**Table S3**. Oligonucleotide primers and synthetic gene fragments.

| **Plasmid** | **Sequence 5'- 3'** | **Description** |
|---|---|---|
| pSC3904 | tccccccccccccctgcaggtcgacCAGGGCAAGTTCGTCGTC | Forward primer to clone upstream region of *rhsF* and *rhsFi* in pKNG101 (Made with Gibson assembly) |
| | taaccaagtcCGCTTTGGAAGCATTGGC | Reverse primer to clone upstream region of *rhsF* and *rhsFi* in pKNG101 (Made with Gibson assembly) |
| | ttccaaagcgGACTTGGTTATAGGCTCGCC | Forward primer to clone downstream region of *rhsF* and *rhsFi* in pKNG101 (Made with Gibson assembly) |
| | tccacggactatagactatactagtGCCAAGCGGGTCGTAGTG | Reverse primer to clone downstream region of *rhsF* and *rhsFi* in pKNG101 (Made with Gibson assembly) |
| pSC3905 | tccccccccccccctgcaggtcgacCAGGGCAAGTTCGTCGTC | Forward primer to clone upstream region of *rhsF* in pKNG101 (Made with Gibson assembly) |
| | taacctcgtaCGCTTTGGAAGCATTGGC | Reverse primer to clone upstream region of *rhsF* in pKNG101 (Made with Gibson assembly) |
| | ttccaaagcgTACGAGGTTATAGAGTAAGGAGTATTTTTATGAG | Forward primer to clone downstream region of *rhsF* in pKNG101 (Made with Gibson assembly) |
| | tccacggactatagactatactagtATGCCAGGGTCTGCTTGG | Reverse primer to clone downstream region of *rhsF* in pKNG101 (Made with Gibson assembly) |
| pSC3911 | GATCAT CTCGAG ACGGGGAAACAGTTCACAGGG | Forward primer to clone sequence of CT domain of *rhsF* and *rhsFi* in pET15-TEV for protein purification (XhoI) |
| | GATCAT GGATCC CTAAGCCCATCTAGATAATG | Reverse primer to clone sequence of CT domain of *rhsF* and *rhsFr* in pET15-TEV for protein purification (XhoI) |
| pSC3912 | GATCAT CCATGG GCAGCAGCCATCATCATCATC | Forward primer to amplify His6Tag-TEV-*rhsF* from pSC3911 (*NcoI*) |
| | GATCAT GGATCC TTACTCTATAACCTCGTACGAAC | Reverse primer to amplify His6Tag-TEV-*rhsF* from pSC3911 (*BamHI*) |

| | | |
|---|---|---|
| | GATCAT CATATG AGTGAACGACTTGAGGCGATA | Forward primer to amplify *rhsFi* to clone into the second MCS (*NdeI*) |
| | GATCAT GGTACC CTAAGCCCATCTAGATAATGGC | Reverse primer to amplify *rhsFi* to clone into the second MCS (*KpnI*) |
| pSC3913 | GGATCCCCCGGGCTGCAG | Forward primer to amplify pBluescript for subcloning of insert into pKNG101 (*BamHI*) |
| | TCTAGAGCGGCCGCCACC | Reverse primer to amplify pBluescript for subcloning of insert into pKNG101 (*XbaI*) |
| | gcggtggcggccgctctagaGTGCTGCTCACCAGCGGC | Forward primer to amplify *vgrG3* fused to HA at C-terminal domain. Insert 1 for Gibson assembly (*XbaI*) |
| | *agcgtaatctggaacatcgtatgggta*GAACAGTTTGGGCAGG<br>CTGG | Reverse primer to amplify *vgrG3* fused to HA at C-terminal domain. Insert 1 for Gibson assembly |
| | *tacccatacgatgttccagattacgct*TGACATCACAAAATTAA<br>AATCCATAAAGTTGAATTGCCATG | Forward primer to amplify *vgrG3* fused to HA at C-terminal domain. Insert 2 for Gibson assembly |
| | tcctgcagcccgggggatccCTGTGCCAGTGCTGCGCA | Reverse primer to amplify *vgrG3* fused to HA at C-terminal domain. Insert 2 for Gibson assembly (*BamHI*) |

| **Plasmid** | **Details of synthetic insert** |
|---|---|
| pSC3948 | Synthetic insert containing coding sequences for RhsF-CT tagged with 3×FLAG at the N-terminus, and RhsFi. |
| | CAGAGGAATACATATATGGACTACAAAGACCATGACGGTGATTATAAAGATCATGATATCGAT<br>TACAAGGATGACGATGACAAAACGGGGAAACAGTTCACAGGGACGGTTTACCGGGCACTTACA<br>CCGAAGCAAAAGCAATGTGCATTAAAAAATCAAGATATATCCCCAAAGGACCCTAGCGCTAGT<br>TATAGCCCTCAGGAACATGTGGAAAATGGAAAACTGCAAACACAGTTTATTTCAACAACTAAA<br>AGTGAAAAAACAAGCGACTTTTATAATAAATGCAATTGTAAAATAAAAGTAGATCTGTCAAAG<br>ATACCTGATTGCGACATCATTGATGTTAGCAAGGGCCAAGGATTGACAGGAAAAGCTAAGCGG<br>TTTGCCACTAAAGACCAAGAAGTATTAATAAGAAATGGCATTCCTAAGGGTTCGTACGAGGTT<br>ATAGAGTAAGTCGACGTAAGGAGTATTTTTATGAGTGAACGACTTGAGGCGATAAAAAAGAAT<br>TTAGATGATTTTTATGATAAATCTTTTGATTCGGATGATATAGAGAGAGCTGAAAATAAAAGTA<br>TCAAAGAAGAGATAGTGGACCTGATAATTCATGCACATAAAAATAGGGATTATCAGCTTGTGA<br>AGGAAAGTATTGATGTTTTGATTGAAAACACAGGATGCCAAGAGGATTTTGAAATTCTTGAAG<br>AAATAGTTTCTCCTCTTCAATCTGCTGGGATTTTGTCGGACCCTGAACTGAATGACTTGGTTATA<br>GGCTCGCCATTATCTAGATGGGCTTAGGTCGAC |
| pSC3949 | Synthetic insert containing coding sequences for RhsF-CT R1403A with 3×FLAG at the N-terminus, and RhsFi. |
| | CAGAGGAATACATATATGGACTACAAAGACCATGACGGTGATTATAAAGATCATGATATCGAT<br>TACAAGGATGACGATGACAAAACGGGGAAACAGTTCACAGGGACGGTTTACGCTGCACTTACA<br>CCGAAGCAAAAGCAATGTGCATTAAAAAATCAAGATATATCCCCAAAGGACCCTAGCGCTAGT<br>TATAGCCCTCAGGAACATGTGGAAAATGGAAAACTGCAAACACAGTTTATTTCAACAACTAAA<br>AGTGAAAAAACAAGCGACTTTTATAATAAATGCAATTGTAAAATAAAAGTAGATCTGTCAAAG<br>ATACCTGATTGCGACATCATTGATGTTAGCAAGGGCCAAGGATTGACAGGAAAAGCTAAGCGG<br>TTTGCCACTAAAGACCAAGAAGTATTAATAAGAAATGGCATTCCTAAGGGTTCGTACGAGGTT |

| | |
|---|---|
| | ATAGAGTAAGTCGACGTAAGGAGTATTTTTATGAGTGAACGACTTGAGGCGATAAAAAAGAAT<br>TTAGATGATTTTTATGATAAATCTTTTGATTCGGATGATATAGAGAGAGCTGAAAATAAAAGTA<br>TCAAAGAAGAGATAGTGGACCTGATAATTCATGCACATAAAAATAGGGATTATCAGCTTGTGA<br>AGGAAAGTATTGATGTTTTGATTGAAAACACAGGATGCCAAGAGGATTTTGAAATTCTTGAAG<br>AAATAGTTTCTCCTCTTCAATCTGCTGGGATTTTGTCGGACCCTGAACTGAATGACTTGGTTATA<br>GGCTCGCCATTATCTAGATGGGCTTAGGTCGA |
| pSC3950 | Synthetic insert containing coding sequences for RhsF-CT Y1456A with 3×FLAG at the N-terminus, and RhsFi. |
| | CAGAGGAATACATATATGGACTACAAAGACCATGACGGTGATTATAAAGATCATGATATCGAT<br>TACAAGGATGACGATGACAAAACGGGGAAACAGTTCACAGGGACGGTTTACCGGGCACTTACA<br>CCGAAGCAAAAGCAATGTGCATTAAAAAATCAAGATATATCCCCAAAGGACCCTAGCGCTAGT<br>TATAGCCCTCAGGAACATGTGGAAAATGGAAAACTGCAAACACAGTTTATTTCAACAACTAAA<br>AGTGAAAAAACAAGCGACTTTgcgAATAAATGCAATTGTAAAATAAAAGTAGATCTGTCAAAGA<br>TACCTGATTGCGACATCATTGATGTTAGCAAGGGCCAAGGATTGACAGGAAAAGCTAAGCGGT<br>TTGCCACTAAAGACCAAGAAGTATTAATAAGAAATGGCATTCCTAAGGGTTCGTACGAGGTTA<br>TAGAGTAAGTCGACGTAAGGAGTATTTTTATGAGTGAACGACTTGAGGCGATAAAAAAGAATT<br>TAGATGATTTTTATGATAAATCTTTTGATTCGGATGATATAGAGAGAGCTGAAAATAAAAGTAT<br>CAAAGAAGAGATAGTGGACCTGATAATTCATGCACATAAAAATAGGGATTATCAGCTTGTGAA<br>GGAAAGTATTGATGTTTTGATTGAAAACACAGGATGCCAAGAGGATTTTGAAATTCTTGAAGA<br>AATAGTTTCTCCTCTTCAATCTGCTGGGATTTTGTCGGACCCTGAACTGAATGACTTGGTTATAG<br>GCTCGCCATTATCTAGATGGGCTTAGGTCGA |
| pSC3951 | Synthetic insert containing coding sequences for RhsF-CT E1497A with 3×FLAG at the N-terminus, and RhsFi |
| | CAGAGGAATACATATATGGACTACAAAGACCATGACGGTGATTATAAAGATCATGATATCGAT<br>TACAAGGATGACGATGACAAAACGGGGAAACAGTTCACAGGGACGGTTTACCGGGCACTTACA<br>CCGAAGCAAAAGCAATGTGCATTAAAAAATCAAGATATATCCCCAAAGGACCCTAGCGCTAGT<br>TATAGCCCTCAGGAACATGTGGAAAATGGAAAACTGCAAACACAGTTTATTTCAACAACTAAA<br>AGTGAAAAAACAAGCGACTTTTATAATAAATGCAATTGTAAAATAAAAGTAGATCTGTCAAAG<br>ATACCTGATTGCGACATCATTGATGTTAGCAAGGGCCAAGGATTGACAGGAAAAGCTAAGCGG<br>TTTGCCACTAAAGACCAAgcgGTATTAATAAGAAATGGCATTCCTAAGGGTTCGTACGAGGTTAT<br>AGAGTAAGTCGACGTAAGGAGTATTTTTATGAGTGAACGACTTGAGGCGATAAAAAAGAATTT<br>AGATGATTTTTATGATAAATCTTTTGATTCGGATGATATAGAGAGAGCTGAAAATAAAAGTATC<br>AAAGAAGAGATAGTGGACCTGATAATTCATGCACATAAAAATAGGGATTATCAGCTTGTGAAG<br>GAAAGTATTGATGTTTTGATTGAAAACACAGGATGCCAAGAGGATTTTGAAATTCTTGAAGAA<br>ATAGTTTCTCCTCTTCAATCTGCTGGGATTTTGTCGGACCCTGAACTGAATGACTTGGTTATAGG<br>CTCGCCATTATCTAGATGGGCTTAGGTCGA |
